# Dynamic task-linked switching between brain networks - A Tri-Network perspective

**DOI:** 10.1101/2020.10.19.344101

**Authors:** Saurabh Bhaskar Shaw, Margaret C. McKinnon, Jennifer Heisz, Suzanna Becker

## Abstract

The highly influential tri-network model proposed by Menon integrates 3 key intrinsic brain networks — the central executive network (CEN), salience network (SN), and the default mode network (DMN), into a single cohesive model underlying normal behaviour and cognition. A large body of evidence suggests that abnormal intra- and inter- network connectivity between these three networks underlies the various behavioural and cognitive dysfunctions observed in patients with neuropsychiatric conditions such as PTSD and depression. An important prediction of the tri-network model is that the DMN and CEN networks are anti-correlated under the control of the SN, such that if a task engages one of the two, the SN inhibits the activation of the other. To date most of the evidence surrounding the functions of these three core networks comes from either resting state analyses or in the context of a single task with respect to rest. Few studies have investigated multiple tasks simultaneously or characterized the dynamics of task switching. Hence, a careful investigation of the temporal dynamics of network activity during task switching is warranted. To accomplish this we collected fMRI data from 14 participants that dynamically switched between a 2-back working memory task and an autobiographical memory retrieval task, designed to activate the CEN, DMN and the SN. The fMRI data were used to 1. identify nodes and sub-networks within the three major networks involved in task-linked dynamic network switching, 2. characterize the temporal pattern of activation of these nodes and sub-networks, and finally 3. investigate the causal influence that these nodes and sub-networks exerted on each other. Using a combination of multivariate neuroimaging analyses, timecourse analyses and multivariate Granger causality measures to study the tri-network dynamics, the current study found that the SN co-activates with the task-relevant network, providing a mechanistic insight into SN-mediated network selection in the context of explicit tasks. Our findings also indicate active involvement of the posterior insula and some medial temporal nodes in task-linked functions of the SN and DMN, warranting their inclusion as network nodes in future studies of the tri-network model. These results add to the growing body of evidence showing the complex interplay of CEN, DMN and SN nodes and sub-networks required for adequate task-switching, and characterizes a normative pattern of task-linked network dynamics within the context of Menon’s tri-network model.

## 1 Introduction

Traditionally the functional correlates of brain activity have been studied one region at a time. However, mounting evidence indicates that the brain is organized into functional networks of interacting brain regions. These clusters of sub-regions fire in a functionally correlated and synchronous manner, and are engaged for specific cognitive tasks, and are hence called functional networks (FN). Out of a large number of previously identified FNs (Thomas Yeo *et al*., 2011) (some of which are shown in Figure 1), three major brain networks are of particular interest to the cognitive neuroscience community. The central executive network (CEN), salience network (SN) and default mode network (DMN) form a tri-network system that is postulated to explain much of cognition and behaviour (Menon, 2011). The CEN is responsible for externally-directed cognitive tasks including working memory and other executive functions, while the DMN is involved in internally-directed cognitive tasks including autobiographical memory retrieval, imagining the future, spatial planning and navigation and self-reflection. Even while the person is not outwardly performing any explicit task (i.e. resting state), the brain is still active with internally-directed cognitive activities driven by the DMN, such as daydreaming, mind-wandering or more focused thought. According to Menon’s tri-network model, the SN is involved in modulating the switching of attention between cognitive processes subserved by the CEN and DMN. This mediates the switch between externally-directed and internally-directed thoughts, respectively, a process required for successful cognition and balanced emotion regulation. Importantly, the inter- and intra-network connectivity between these three core networks is found to be dysregulated in psychiatric conditions such as depression (Hamilton *et al*., 2011; Yang et al., 2016), schizophrenia (Moran *et al*., 2013) and PTSD (Menon, 2011). Hence, studying the tri-network activity and its switching dynamics is essential to understanding the neurological underpinnings of the aberrant thought processes and emotional dysregulation observed in such complex neuropsychological disorders. This study investigates the temporal dynamics of various sub-components of the tri-network model with a composite memory task designed to activate the three networks.

**Figure 1:**
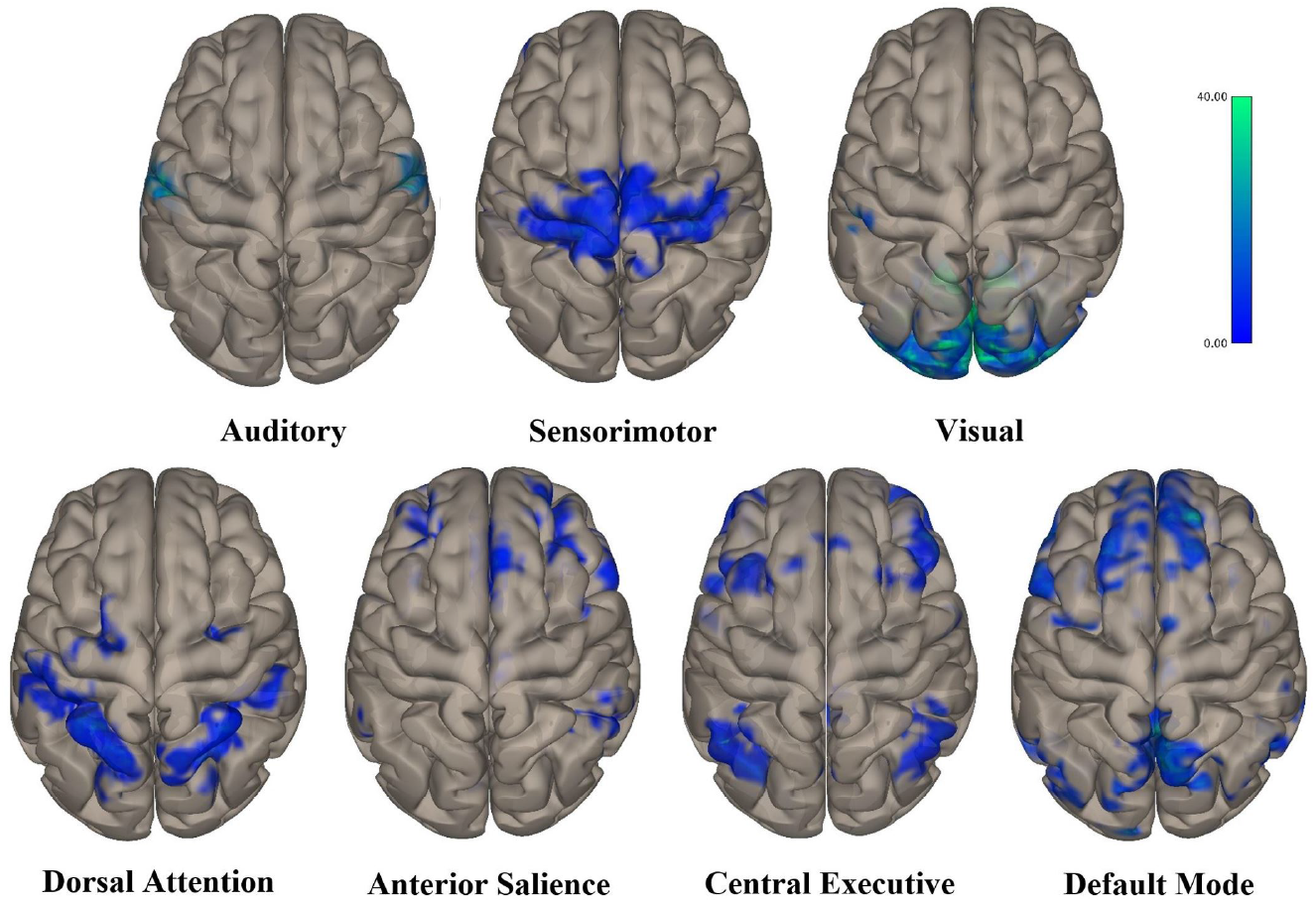
Seven major functional networks (FN) found using spatiotemporal ICA analyses on fMRI BOLD signal, labeled based on results from Shirer et al. (2012). Activity shown is thresholded at FDR-corrected p < 0.05.

**Figure 2:**
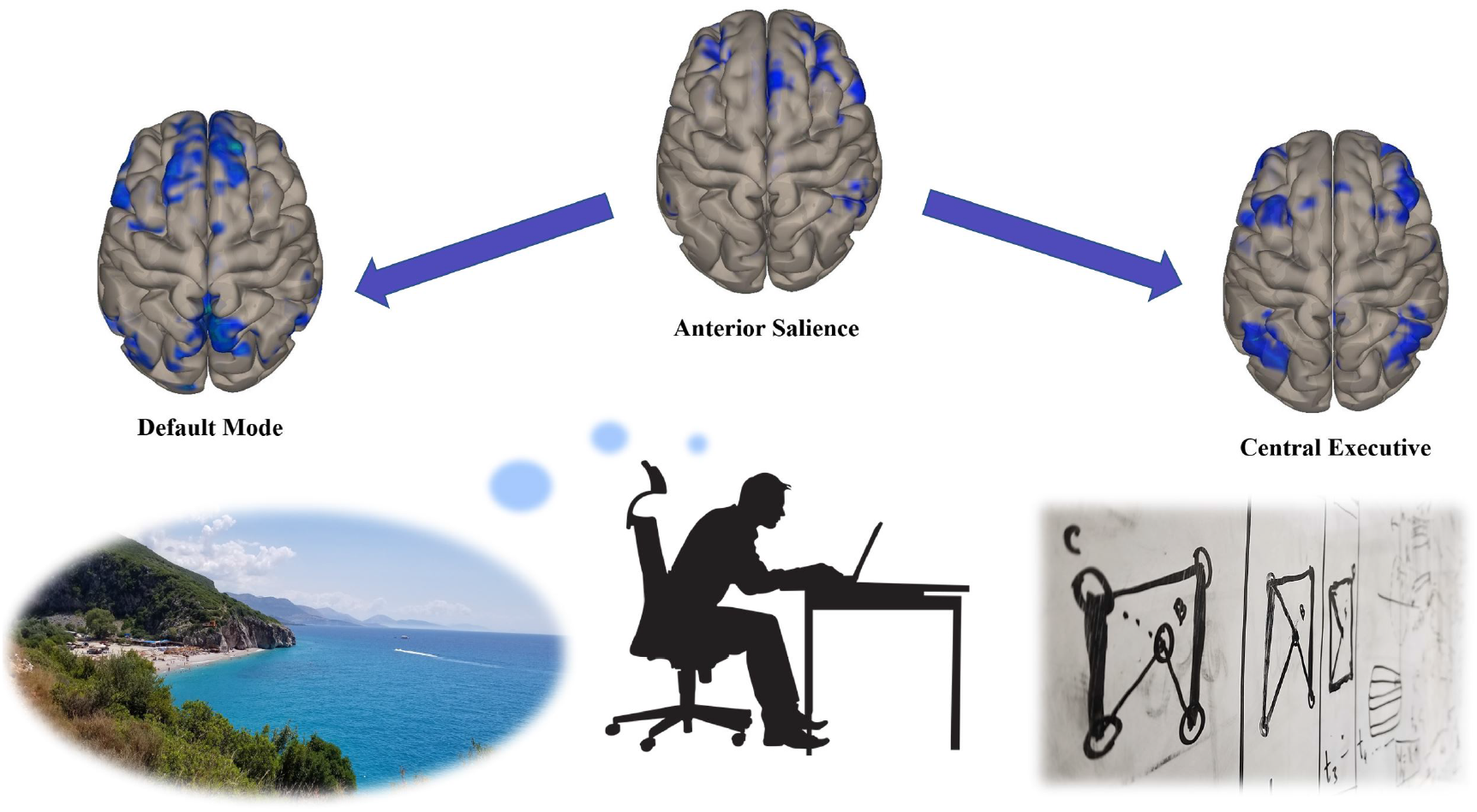
The Tri-Network model, as described by Menon (2011), postulates that appropriate processing of everyday tasks requires switching between self-related or interoceptive processing and externally directed or exteroceptive processing reliant on the default mode network (DMN) and the central executive network (CEN) respectively. The switching between these networks is regulated by the salience network (SN), as show in this schematic.

### 1.1 The Tri-Network Model

An important prediction of the tri-network model is that the DMN and CEN are anticorrelated under the control of the SN, such that if a task engages one of the two, the SN inhibits the activation of the other. To date most of the evidence surrounding the functions of these three core networks comes from either resting state analyses (Goulden *et al*., 2014; Taghia *et al*., 2017; Ryali *et al*., 2016) or paradigms where brain network activation is studied in the context of a single task relative to rest (Sidlauskaite *et al*., 2014). Few studies have investigated multiple tasks simultaneously (see Spreng *et al*. (2010)), and even fewer have attempted to characterize the dynamics of task switching(see Bréchet *et al*. (2019); Sidlauskaite *et al*. (2014)). Bréchet *et al*. (2019) characterized the spatial regions activated across two different tasks, while describing the temporal dynamics using EEG microstates, a method that makes some faulty assumptions (Shaw *et al*., 2019). Hence, a detailed investigation of the temporal dynamics of network activity during task switching is warranted.

Next, we briefly review the prominent nodes and subnetworks within the DMN, CEN and SN and their involvement in network switching, as prescribed by the tri-network model.

#### 1.1.1 Default Mode Network (DMN)

The default mode network (DMN) consists of some key nodes within the pre-frontal cortex (PFC), posterior parietal cortex (PPC) and medial temporal lobe (MTL) (Raichle, 2015) and is the predominantly active network when the person is not performing any particular task (Raichle, 2015). For this reason, this network was initially mis-characterized as simply reflecting the brain’s idling state and was also referred to as the default network or task-negative network. However, a large body of evidence shows that the DMN subserves a wide range of cognitive tasks involved in internally-directed thoughts, spanning numerous distinct functional domains, which map onto three distinct sub-networks (Andrews-Hanna *et al*., 2014; Thomas Yeo *et al*., 2011). These are

- the dorso-medial sub-network comprised of nodes in the lateral temporo-parietal cortices involved in attention and some aspects of social cognition functions (Krall *et al*., 2015);
- the medio-temporal sub-network, primarily consisting of nodes within the medial temporal lobe and posterior parietal cortex subserving self-referential memory functions such as autobiographical memory retrieval, episodic future thinking and contextual memory retrieval;
- and the core network composed of cortical-midline structures, such as the posterior cingulate cortex (PCC) and antero-medial prefrontal cortex (amPFC), responsible for integrating information across the other two subnetworks.

Self-referential memory encoding and retrieval processes were originally thought to be left and right lateralized respectively (Craik *et al*., 1999), however, there is now considerable evidence for the left lateralization of self-referential processing (Axelrod *et al*., 2017) and autobiographical memory retrieval (Kim, 2012), with increasingly bilateral activity in individuals with highly superior autobiographical memory (Mazzoni *et al*., 2019).

#### 1.1.2 Central Executive Network (CEN)

The central executive network (CEN) is a complex attentional control system that is anchored in the dorso-lateral prefrontal cortex (dlPFC) and some parietal regions. For this reason, it is sometimes referred to as the frontoparietal attention, frontoparietal control network, or task-positive network and is responsible for a host of executive functions such as updating working memory, inhibitory control, selective attention, multiple task coordination (Collette and Van der Linden, 2002; Baddeley, 1996), and random sequence generation (Baddeley *et al*., 1998).

Executive functions of the CEN require integration of information from different sensory modalities to create a modality-invariant representation of items in working memory. To this end, two prominent subsystems feed into the dlPFC, namely the visuospatial sketchpad and the phonological loop that instantiate and maintain visuospatial and verbal object representations respectively (Baddeley, 2000, 2003). These distributed patterns of modality-specific information are integrated at parietal and prefrontal structures over increasing time lags (Christophel *et al*., 2017).

#### 1.1.3 Salience Network (SN)

The salience network is implicated in detecting the behavioural, emotional and reward-linked salience of stimuli and in directing attention toward those stimuli. It is primarily composed of two major cortical nodes, the anterior insula (AI) and the dorsal anterior cingulate cortex (dACC) (Menon, 2011). The AI integrates information across sensory afferents to detect stimuli coherent with behaviourally relevant actions in a goal-directed manner, communicating this to the dACC, which uses this information in directing attention and response selection (Sridharan *et al*., 2008; Menon and Uddin, 2010; Menon, 2015). These responses are in turn communicated to somatomotor areas, hypothalamus and the periaqueductal grey (PAG) via efferents. The dACC also has some reciprocal connections to the AI, providing it with interoceptive and autonomic information.

Using the emotional, social or task-linked salience of an input, the SN is also thought to act as a “gate” by activating the relevant functional network to further process and respond with an appropriate action (Menon, 2015), as described in the following section.

### 1.2 Dynamic Interaction amongst Intrinsic Networks

Under Menon’s 2011 tri-network model, the SN acts as a gate by activating the DMN for internally-directed and self-relevant tasks, and the CEN for externally-directed executive tasks.

There is a large body of work, using dynamic causal modelling and Granger causality analysis, supporting SN’s gating function between major network nodes (Friston *et al*., 2014) during resting state (Sridharan *et al*., 2008; Goulden *et al*., 2014; Chand *et al*., 2017), visual or auditory attention-switching (Sridharan *et al*., 2008), Go-No-Go and congruent/incongruent flanker tasks (Cai *et al*., 2016) and magnitude and parity judgement tasks (Sidlauskaite *et al*., 2014). These causal analyses support the gating function of the SN within tasks (Chand and Dhamala, 2016) and even more so when switching between externally-oriented tasks (Sidlauskaite *et al*., 2014).

These studies identify the right Anterior Insula (rAI) as the node with the lowest latency of the event-related activity and the highest net causal ‘‘outflow” to other SN, CEN and DMN nodes, implicating a central role for rAI as a hub responsible for switching between different tasks, requiring a switch in the predominantly active network (Sridharan *et al*., 2008). Neurofeedback training is also known to improve AI-linked network regulation of the CEN and DMN, supporting the hub-like role of this region (Zhang *et al*., 2015). Furthermore, stimulating other cortical nodes of the salience and central executive networks using transcranial magnetic stimulation (TMS) causes deactivation of DMN nodes, suggesting that it is under the inhibitory control of CEN/SN nodes (Chen *et al*., 2013).

However, most of the work surrounding the gating function of SN has either been in the context of resting state (Chand *et al*., 2017), or in the context of suppressing DMN and recruiting CEN during externally-directed tasks (Sridharan *et al*., 2008; Chen *et al*., 2013; Chand and Dhamala, 2016), and has not assessed the network switching dynamics when recruiting DMN for self-related tasks in healthy participants. Furthermore, these studies do not look at the temporal sequence of activity while switching between tasks that rely on different networks, that can provide a dynamic view of the switching process.

This study attempts to address these shortcomings by studying three major aspects of CEN, SN and DMN functioning in the context of a paradigm that involves dynamic task switching designed to activate these three networks. This involves

1. identification of the nodes and sub-networks within the three major networks that are involved in such dynamic network switching,
2. characterization of the temporal pattern of activation of these nodes and sub-networks, and finally
3. investigation of the causal influence that these nodes and sub-networks exert on each other over the course of the tasks.

To fulfill these aims, the spatiotemporal pattern of activating each of the networks while performing the tasks is first described using data-driven patterns of voxel-to-voxel connectivity and event-related activity timecourses, followed by multivariate Granger causality analysis to probe the patterns of causality between the network nodes.

## 2 Methods

In this study, fMRI data were collected from 14 healthy participants (8 females, mean age 22±3 years) while they performed a complex memory task expected to evoke dynamic switching between intrinsic networks. The complex memory task consisted of randomly interleaved blocks of autobiographical memory (ABM) retrieval trials and working memory (WM) trials, expected to activate the DMN and CEN respectively. At the end of some blocks, a task-switching cue prepared participants to switch from an AM to a WM block or vice versa, and was expected to activate the SN. All participants provided informed consent prior to their participation in the study, in accordance with the approved Hamilton Integrated Research Ethics Board (HiREB) protocol.

### 2.1 Memory Assessment

Each participant engaged in a pre-scanning phase, during which detailed autobiographical memories were recorded by the participant. Following this, participants completed a 1 hour 20 min long memory assessment in the MRI scanner, comprised of randomly ordered 30-second blocks of either cued AM retrieval (remembering previous autobiographical memories - ABM) or a 2-back WM task (remembering which word they saw 2 words ago in a stream of words). These two block types were predicted to activate the DMN and CEN respectively, while a cue to a pending task switch between blocks was expected to activate the SN. Participants completed as many blocks as possible within 80 minutes, up to a maximum of 64 blocks (32 ABM, 32 WM), in chunks of 16 blocks. Each block was followed by a 60 second rest period to prevent fatigue.

#### Autobiographical memory (ABM) task

Our ABM task was modeled after a similar protocol used by Addis *et al*. (2004), with some modifications to balance the visual stimulus processing and motor response demands between the ABM and WM tasks. Prior to entering the scanner, the participant was asked to recall and record within an Excel sheet up to 10 positive or neutral autobiographical memories in vivid detail. The participant was then asked to identify descriptive words corresponding to each memory, that would serve as cues during the ABM retrieval task. During each ABM block within the scanner, the participant was shown a series of cue words pertaining to one of the previously described autobiographical memories and was instructed to vividly imagine the corresponding memory in detail. Each ABM block included 10 cues pertaining to the same memory, shown for 2 seconds each. In each different ABM block, a different memory was cued. To ensure that the participant was staying on task, at random points within each block, they were asked to perform a 4-alternative forced choice trial in which they should select the word that represents the memory they were currently recalling from a selection of four words.

#### Working Memory (WM) task

During the WM blocks, the participant was presented with a sequence of neutral words (not expected to elicit the autobiographical memories recorded) and was asked to remember the word that was presented two words back. At random points within each block, they were asked to perform a 4-alternative forced choice trial in which they should choose the word they saw two words ago from a selection of four words.

#### Task cue period

The words Word Memory and Autobiographical Memory were shown for 2 seconds before the onset of the WM and ABM blocks, respectively. A schematic illustrating the task paradigm is shown in Figure 3.

**Figure 3:**
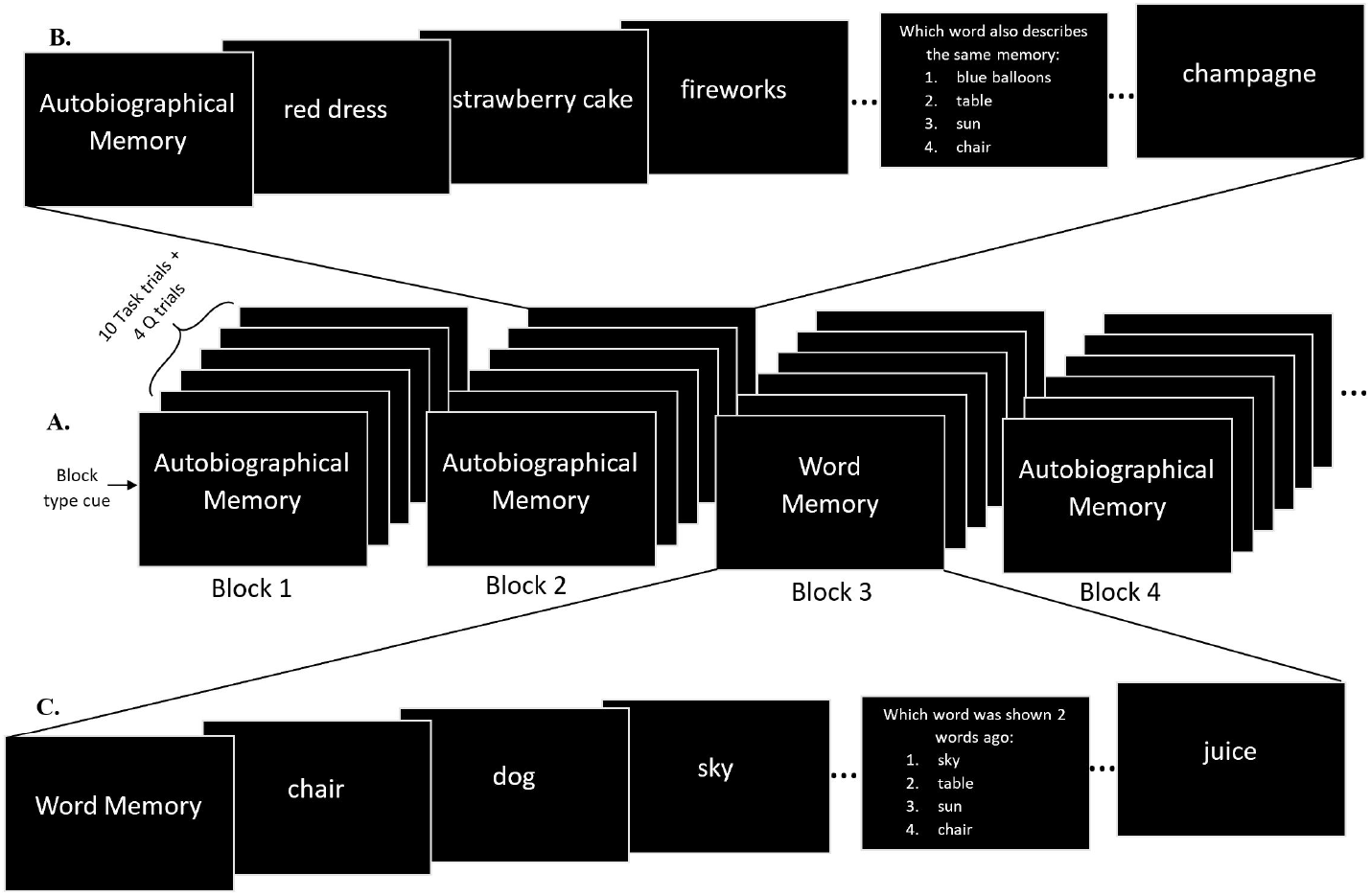
A. Schematic of the dynamic switching task. Each block is either an autobiographical memory retrieval task (ABM task) or a 2-back working memory (WM) task. Each block consists of exactly 10 task cue trials and 4 four alternative forced choice (4-AFC) trials placed at random locations within the block. Each trial is presented to the participant for 2 seconds, except the 4-AFC trials that persist until the participant has made a choice. The structure of an ABM block and a WM block are shown in more detail in panels B (top) and C (bottom) respectively. During the ABM blocks (panel B), the task cues are memory keywords that correspond to one of the participant’s autobiographical memories, thereby cuing that particular memory. The trials within a block consists of different memory cues from a single autobiographical memory. On the other hand, the task cues during the WM blocks (panel C) are a sequence of words from which the participant needs to remember the word shown 2 words ago.

### 2.2 MRI acquisition

A GE MR750 3T MRI scanner and an 8 channel RF coil (General Electric Healthcare, Milwaukee, WI) were used for data acquisition. FMRI data were acquired using an axial 3D fSPGR pulse sequence (2D gradient echo EPI, FA=90°, TE/TR=35/2000ms, 64×64 matrix, 39 interleaved 3.8 mm slices, 1200 temporal points).

### 2.3 MRI data analysis

The raw fMRI data were first bandpass filtered (0.008Hz 0.09Hz), motion corrected, and aligned with the MNI standard space after applying a 2mm Gaussian blur. These preprocessed fMRI data were used for all subsequent analyses.

To accomplish the three major study aims described at the end of section 1, we conducted a series of analyses probing the spatio-temporal patterns of activity in the three networks of interest. These analyses are discussed below, in the same order as the corresponding study objectives they are accomplishing.

1. We first identified brain regions differentially activated by the ABM and WM tasks, and during task switching, using whole-brain connectome multivariate pattern analysis (connectome-MVPA), followed by a group-wide independent component analysis (group-ICA). Both of these analyses are data-driven and hence avoided any a-priori biases or assumptions around the network nodes and sub-networks that might participate in the tasks being studied. The group-ICA components corresponding to independent CEN, DMN and SN sub-networks were then identified for further analysis.
2. Next, temporal patterns of task-linked network activation were studied using two complementary approaches. First, an ICA-based data-driven approach was used to identify and study the temporal activation patterns of the three networks. This was followed by a region of interest (ROI)-driven analysis, based on evidence from the literature to identify temporal patterns of activity within key nodes of the CEN, DMN and SN.
3. Finally, these activity time courses were analyzed using multivariate Granger causality (MVGC) analysis to identify the causal influence of these nodes and sub-networks on one another.

These analyses were carried out using the SPM12 and CONN toolboxes (Whitfield-Gabrieli and Nieto-Castanon, 2012a), along with custom MATLAB scripts and are described in detail in the following sub-sections.

#### 2.3.1 Connectome Multivariate Pattern Analysis (connectome-MVPA)

In order to identify brain regions activated by the ABM and WM tasks without any a-priori biases (study aim 1), we used data-driven whole-brain connectome multivariate pattern analysis (connectome-MVPA). This analysis identifies whole-brain multivariate patterns of functional connectivity (*MCOR*), allowing for discovery of task-linked patterns of voxel-to-voxel functional connectivity (Beaty *et al*., 2015; Thompson *et al*., 2015; Whitfield-Gabrieli *et al*., 2016).

Mathematically, the *MCOR* connectivity maps refer to the top *M* spatial principal component scores of the seed-based correlation matrix containing the correlation between each seed voxel and all other voxels in the brain, aggregated across all participants and conditions.

The whole-brain connectome-MVPA maps are identified by repeating the following procedure for each of the *K* voxels, *x*, in the brain. First, for each voxel *x*, the 1 × (*K*)-dimensional seed-based correlation map is computed using *x* as the seed, for each participant-condition pair. Assuming a total of *N* participant-condition pairs, these seedbased correlation maps are aggregated into a *N × K* seed-based correlation matrix (**R**) for each voxel *x*.

Next, the top *M* principal spatial components of **R** are identified using principal component analysis (PCA), as shown in equation 1.

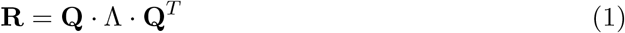

where Λ is the *M × M* square diagonal matrix of eigenvalues corresponding to the principal spatial components (eigenvectors) contained in the *K × M* dimensional **Q** matrix. These components form an orthogonal basis set that span the space of the seed-based correlation patterns observed across all participants and conditions.

The *N × M* dimensional component scores matrix (MCOR) is computed using the **R** matrix and **Q** as shown in Equation 2, giving a 1 × *M* dimensional component score for each of the *N* participant-condition pairs, for each voxel *x*:

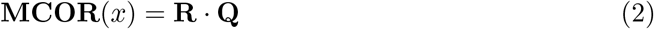

Consequently, each element *r*(*x,y*), of the *n^th^* participant-condition pair seed-based correlation matrix (**R**) can be written as,

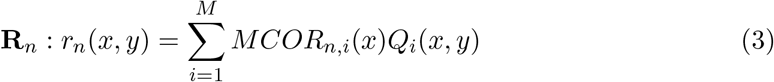

The *MCOR* maps provide a multivariate low-dimensional representation of the between-subjects/conditions variance of the seed-based functional connectivity between each voxel *x* and the rest of the brain. Task-linked differences in these whole-brain patterns of connectivity can then be identified by performing a second-level between condition/participant analysis by performing an F-contrast on the *MCOR* spatial maps between the task conditions of interest.

First-level connectome-MVPA analyses were run using the *CONN* toolbox (Whitfield-Gabrieli and Nieto-Castanon, 2012b) with *M* = 4 top components, followed by a second-level contrast between the ABM and WM task conditions to identify the multivariate voxel-to-voxel patterns of connectivity that differ between these task conditions.

#### 2.3.2 Independent Component Analysis (group-ICA)

We studied the temporal dynamics of the 3 networks (study aim 2) using two different approaches, a data-driven approach to identifying the three networks and a more traditional ROI-driven approach. For the data-driven approach, following connectome-MVPA, we used group wide independent component analysis (group-ICA) for the identification of spatio-temporally independent CEN, DMN and SN sub-networks without any a-priori biases (study aim 1).

The group-ICA decomposition (Calhoun *et al*., 2009) was performed on the pre-processed voxel data using *tanh* as the non-linear contrast function with the iterative FastICA algorithm to identify 20 group-ICA spatial components. The group-level ICA components were labeled based on their spatial overlap (quantified using the Dice coefficient) with known functional networks (described in Shirer *et al*. (2012) and downloaded from https://findlab.stanford.edu/functional_ROIs.html) and back-projected to individual participants’ data, using GICA3 back-projection (Erhardt *et al*., 2011), to obtain the activity timeseries of each ICA component. These timeseries were used for the temporal analysis of CEN, DMN and SN sub-network activity (study aim 2).

#### 2.3.3 Regions-of-Interest (ROI) Definitions

The temporal dynamics of CEN, DMN and SN node activity (study aim 2) were also studied using an ROI approach, based on the average BOLD activity within major nodes of the three networks. Two network configurations were studied, each using a different set of nodes to define the three networks, to identify the appropriate collection of nodes that adequately described tri-network dynamics in the context of a task paradigm. The first network configuration included an abridged set of 10 nodes that has been extensively used to define the three networks in most prior studies investigating the tri-network model. Additionally, since this widely used set of tri-network nodes did not include key DMN and SN nodes, such as medial-temporal lobe (MTL) nodes and the posterior insula (PI), a second network configuration was defined by adding 8 nodes that span the MTL and PI, to the earlier set of nodes comprising the DMN and the SN respectively.

The nodes for the abridged and extended network configurations were defined using a combination of standard and custom regions of interest (ROI), as described in Tables 1 and 2, respectively. The standard ROIs from the Harvard-Oxford atlas (H-O atlas) were used to define the posterior cingulate cortex (PCC), precuneus (PreC), right and left posterior parietal cortex (r/l PPC), hippocampal (r/l HC) and parahippocampal (r/l pHC) ROIs. The rest of the ROIs were created using a sphere of radius 5mm, centered at pre-defined MNI coordinates, given in Tables 1 and 2.

**Table 1:** Definitions of the abridged set of regions of interest (ROI) used to define the three networks for ROI analysis. All ROIs are defined as mm spheres centered at the specified MNI coordinates. The coordinates for prefrontal cortex nodes (vmPFC, amPFC, r/l dlPFC) are based on those reported in Watanabe *et al*. (2013), that for r/l AI are based on Menon (2015), while those for dACC is based on Cai *et al*. (2016). The rest of the ROIs use definitions from the Harvard-Oxford atlas (H-O atlas).

| Network | ROI Name | Abbrev-<br>-iation | MNI Coordinates<br>(x,y,z) mm |
| --- | --- | --- | --- |
| Salience<br>Network | Right Anterior Insula | r AI | (36,18,4) |
|  | Left Anterior Insula | l AI | (-36,18,4) |
|  | Dorsal Anterior Cingulate Cortex | dACC | (7,18,33) |
| Default<br>Mode<br>Network | Ventromedial Prefrontal Cortex | vmPFC | (-3,40,0) |
|  | Posterior Cingulate Cortex | PCC | H-O atlas |
|  | Anteromedial Prefrontal Cortex | amPFC | (1,55,26) |
| Central<br>Executive<br>Network | Right Posterior Parietal Cortex | r PPC | H-O atlas |
|  | Left Posterior Parietal Cortex | l PPC | H-O atlas |
|  | Left Dorsolateral Prefrontal Cortex | l dlPFC | (43,21,38) |
|  | Right Dorsolateral Prefrontal Cortex | r dlPFC | (-48,21,38) |

**Table 2:**
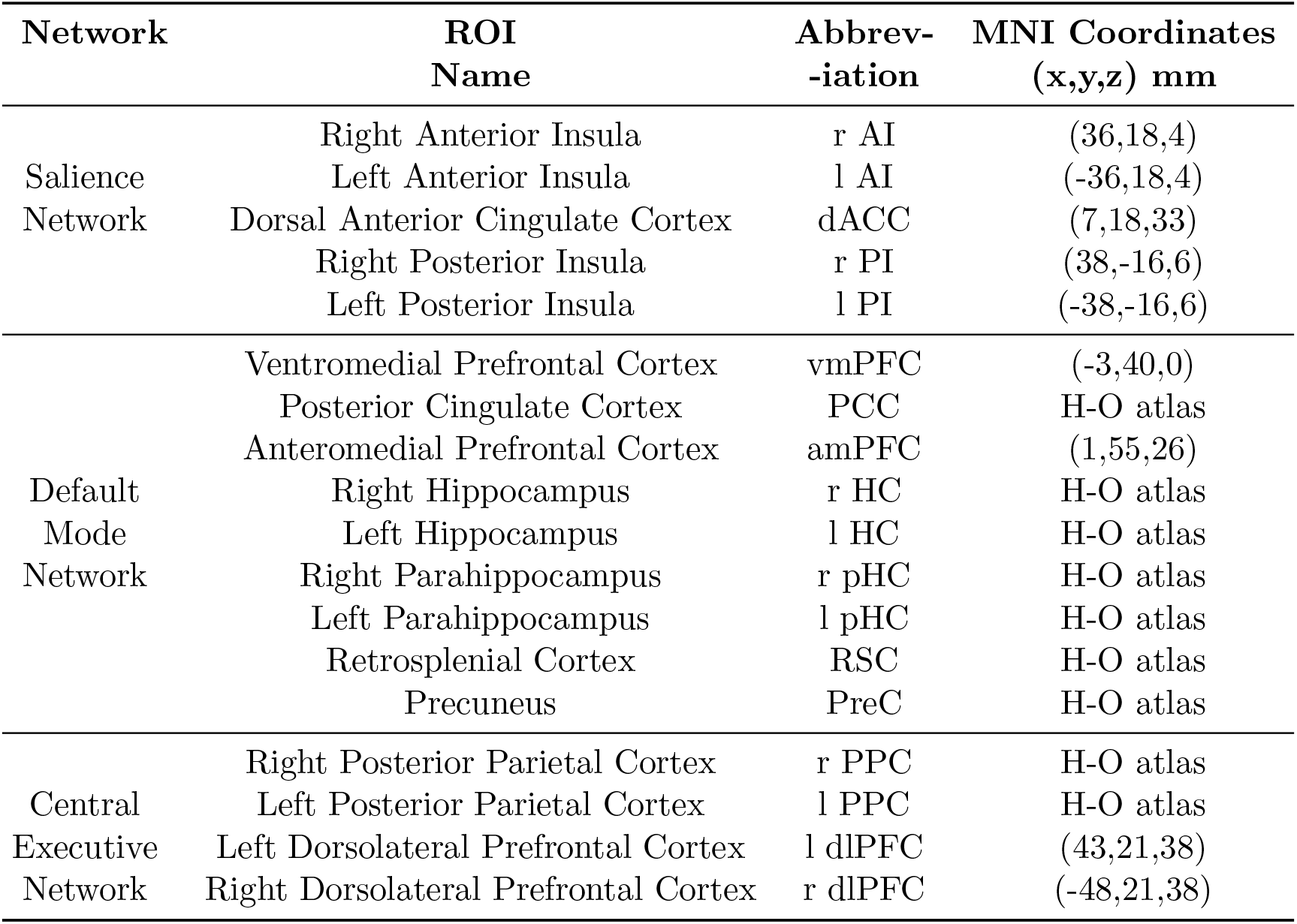
Definitions of the extended set of regions of interest (ROI) used to define the three networks for ROI analysis. All ROIs are defined as mm spheres centered at the specified MNI coordinates. The coordinates for prefrontal cortex nodes (vmPFC, amPFC, r/l dlPFC) are based on those reported in Watanabe *et al*. (2013), that for r/l AI are based on Menon (2015), while those for dACC is based on Cai *et al*. (2016). The definition for r/l PI are based on coordinates reported in Davidovic *et al*. (2019). The rest of the ROIs use definitions from the Harvard-Oxford atlas (H-O atlas).

#### 2.3.4 Multivariate Granger Causality (MVGC) Analyses

In order to assess the causal interactions between the networks (study aim 3), the causal influence between the identified sub-networks and network nodes were studied using multivariate causality analyses on the ICA component timeseries and the ROI timeseries, respectively.

Granger causality has been extensively used to characterize data-driven estimates of directional causality between two sources of activity in EEG and fMRI (Iwabuchi *et al*., 2017; Seth *et al*., 2015). It relies on information-theoretic principles of causality, asserting that a signal *x_i_* causally influences signal *x_j_* if predictions about the future samples of *x_j_* can be improved by including past samples of *x_i_* in addition to past samples of *x_j_* itself. The multivariate version of Granger causality (Barnett and Seth, 2014) extends this univariate description by also considering activity from nodes in the system that may indirectly influence the causality from *i→j*. Within this framework, two vector autoregressive (VAR) models are fitted - a ‘full’ or ‘unrestricted’ VAR model that models the activity of all system nodes (x(*t*)) as a function of their past activity; and a ‘reduced’ or ‘restricted’ VAR model that estimates future activity of the system excluding node *i*(x(*t*)*x_i_*), as shown in Equations 4 and 5 respectively.

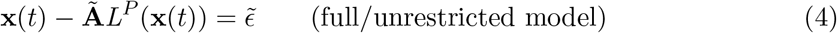

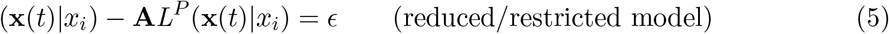

where **A** and 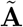 are (*n* − 1) × *P* and *n × P* dimensional coefficient matrices respectively, for a system with n nodes. *L^P^* is the lag operator such that *L^P^*(x(*t*)) gives a vector with the last *P* values in x(*t*), and *ϵ* and 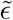 are white time-uncorrelated noise processes with covariance **D** and 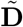 respectively (Duggento *et al*., 2018).

The MVGC strength for the connection from *i→j* is given by estimating the additional information gained about future *x_j_* samples by including previous *x_i_* samples, ignoring the information provided by the other nodes in the system (Equation 6):

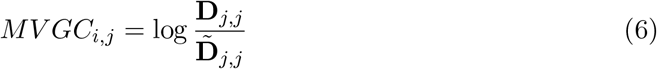

The optimal value for *P* of the lag operator *L^p^* can be estimated from the time series data using the Akaike information criterion (AIC) or Bayesian information criterion (BIC), to maximize the fit of the VAR model while minimizing overfitting (Barnett and Seth, 2014). Optimal values of *P* = 3 have been previously observed when applying MVGC analysis on BOLD fMRI data (Duggento *et al*., 2018).

MVGC analyses were performed on the ICA and ROI timecourses using the MVGC Toolbox (Barnett and Seth, 2014). The AIC was used to estimate the optimal lag for fitting the VAR model, followed by splitting the time courses into 14 second long sliding windows with a step size of 2 seconds (1 TR). This provided a dynamic estimate of the MVGC values over time, that was compared against the the null MVGC estimate using time-permuted surrogate signals. The connections that were significantly different from their surrogate estimates were retained. A unique directed graph was constructed from the MVGC estimates for each sliding window, generating a series of graphs.

To characterize the changing role of each network node in this series of graphs, the net causal outflow was estimated for each network node by subtracting the weighted sum of incoming connections to the node, (i.e. importance weighted in-degree of the node) from the weighted sum of outgoing connections from the node (i.e. importance weighted out-degree of the node). A hubness score was also estimated for each node of these graphs by identifying the nodes that have high node degree, high node betweenness, low clustering coefficients and low average path length. This combination of four sub-measures is thought to indicate hub-like behaviour within a network, and has been used to define a composite hubness score (van den Heuvel *et al*., 2010). This composite hubness score was derived from sub-scores of each of the four sub-measures. A sub-score of 1 was assigned to each node if it ranked in the top 20% of nodes that matched the aforementioned hub-like pattern of the four sub-measures, while a sub-score of 0 was assigned to all nodes that failed to meet this criteria. The sub-scores for each of the four sub-measures were then added together to identify the composite hubness score for each node. Consequently, this score ranged from 0, representing no hub-like behaviour, to 4, indicating the presence of all four hub-like characteristics. Further details on this composite hubness score can be found in van den Heuvel *et al*. (2010).

## 3 Results

This section first discusses the results of the connectome-MVPA and group-ICA analyses in subsection 3.1, accomplishing the goals of study aim 1 to spatially characterize the tri-network activity by identifying sub-networks and dominant regions of interest. Next, the results pertaining to study aim 2 are discussed by exploring the temporal characteristics of the ICA and ROI time courses in subsection 3.2. Finally, the results of the MVGC analyses performed to investigate study aim 3 are presented in subsection 3.3, characterizing the patterns of causality between the identified nodes and sub-networks using multivariate Granger causality analysis.

### 3.1 Spatial characterization of tri-network activity

#### 3.1.1 Connectome-MVPA Analysis

The connectome-MVPA analysis for the contrast between the ABM trials and WM trials revealed that when contrasted against the WM trials, the ABM trials more strongly activated the right and left parietal cortices and the posterior cingulate cortex (PCC), while the WM trials more strongly activated supplementary motor areas, right and left middle frontal gyri, and some medial frontal cortical and right cerebellar regions. These results, shown in Figure 4 and Table 3, are consistent with previous findings of activation within the medio-temporal subnetwork of the DMN in autobiographical memory recall (Andrews-Hanna *et al*., 2014), and activation of phonological loop structures in working memory (Yaple *et al*., 2019).

**Table 3:**
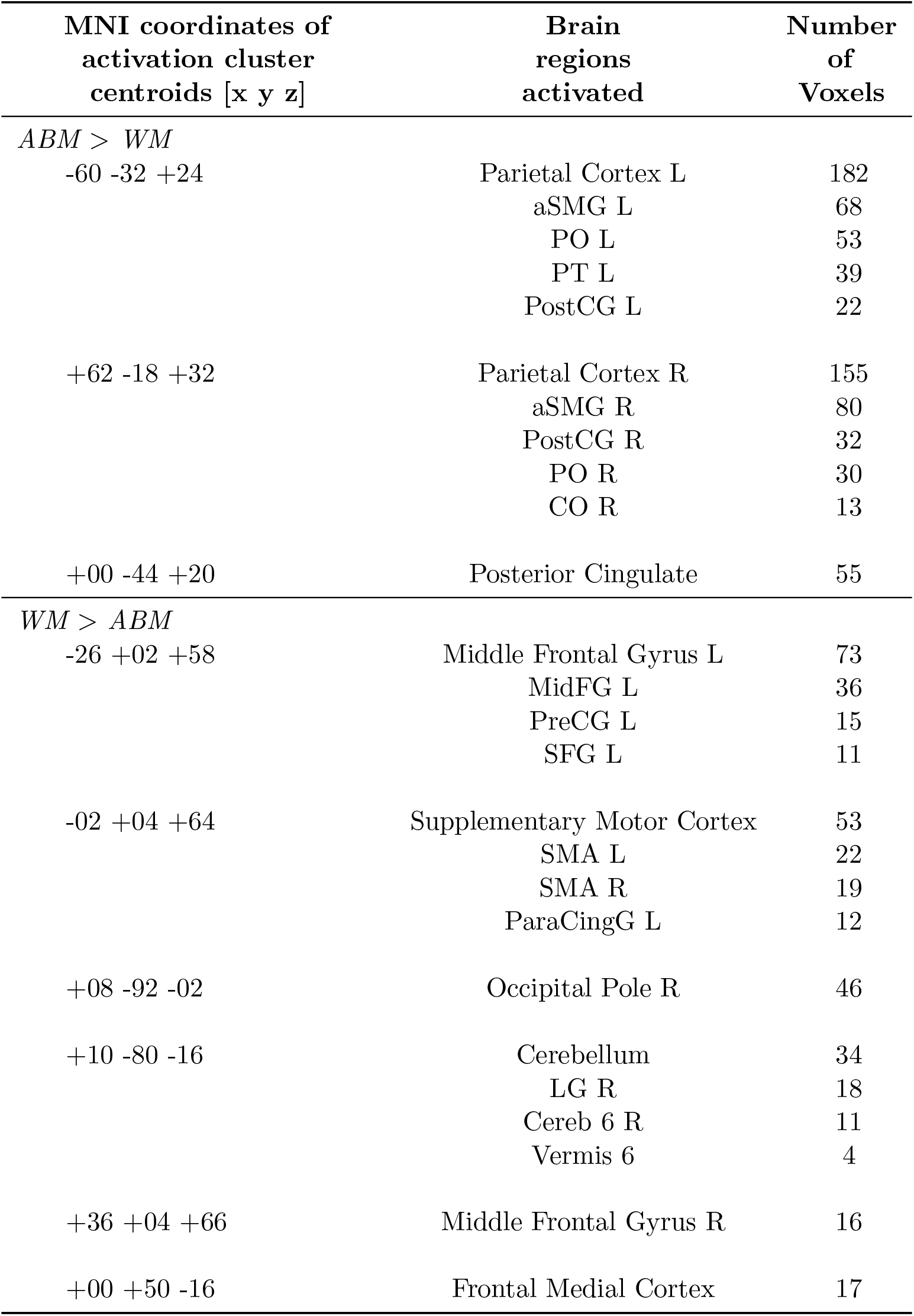
Clusters of voxels identified in a two-sided ABM > WM contrast of connectome-MVPA values. The abbreviations used are aSMG = Anterior Supramarginal Gyrus, PostCG = Post Central Gyrus, PO = Parietal Opercular Cortex, PT = Planum Temporale, CO = Central Opercular Cortex, MidFG = Middle Frontal Gyrus, SFG = Superior Frontal Gyrus, SMA = Supplementary Motor Cortex, LG = Lingual Gyrus.

| MNI coordinates of<br>activation cluster<br>centroids [x y z] | Brain<br>regions<br>activated | Number<br>of<br>Voxels |
| --- | --- | --- |
| *ABM > WM* |  |  |
| -60 -32 +24 | Parietal Cortex L | 182 |
|  | aSMG L | 68 |
|  | PO L | 53 |
|  | PT L | 39 |
|  | PostCG L | 22 |
| +62 -18 +32 | Parietal Cortex R | 155 |
|  | aSMG R | 80 |
|  | PostCG R | 32 |
|  | PO R | 30 |
|  | CO R | 13 |
| +00 -44 +20 | Posterior Cingulate | 55 |
| *WM > ABM* |  |  |
| -26 +02 +58 | Middle Frontal Gyrus L | 73 |
|  | MidFG L | 36 |
|  | PreCG L | 15 |
|  | SFG L | 11 |
| -02 +04 +64 | Supplementary Motor Cortex | 53 |
|  | SMA L | 22 |
|  | SMA R | 19 |
|  | ParaCingG L | 12 |
| +08 -92 -02 | Occipital Pole R | 46 |
| +10 -80 -16 | Cerebellum | 34 |
|  | LG R | 18 |
|  | Cereb 6 R | 11 |
|  | Vermis 6 | 4 |
| +36 +04 +66 | Middle Frontal Gyrus R | 16 |
| +00 +50 -16 | Frontal Medial Cortex | 17 |

**Figure 4:**
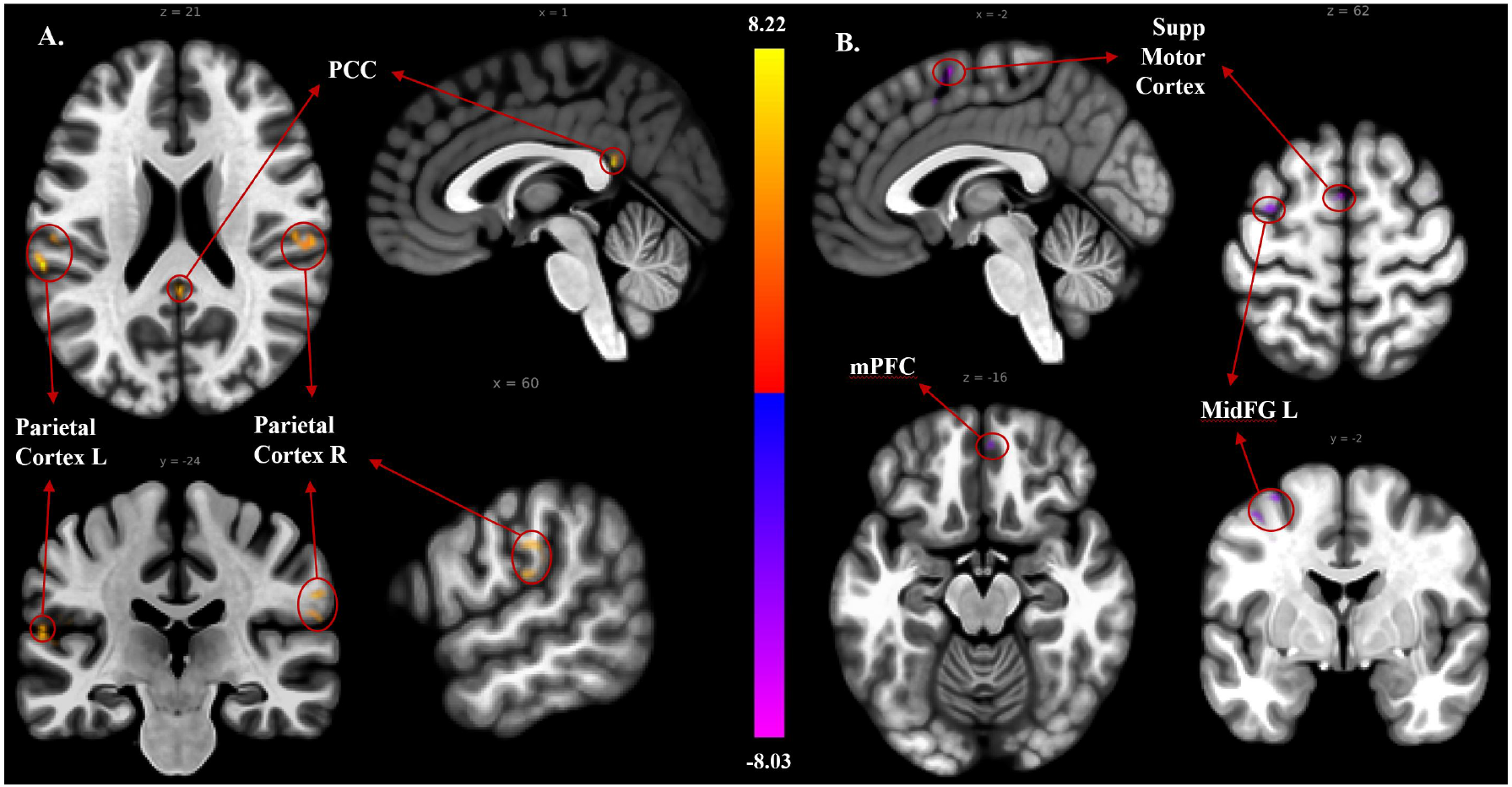
Connectome-MVPA activation for the contrast between ABM>WM with positive contrast clusters (ABM > WM) shown in red/yellow (panel A on the left) and negative contrast clusters (WM > ABM) shown in blue/purple (panel B on the right). The identified clusters of activity, along with their MNI coordinates are listed in Table 3. Abbreviations: PCC - posterior cingulate cortex, mPFC - medial prefrontal cortex, MidFG - Middle Frontal Gyrus, Supp Motor Cortex - Supplementary Motor Cortex. A cluster-discovery threshold of p-uncorrected < 0.001 and cluster-visualization threshold of FDR-corrected p < 0.05 was used to generate this image.

#### 3.1.2 ICA Networks - Spatial Pattern Analysis

Of the 20 identified ICA components, 11 were found to have significant overlap with various functional networks (Shirer *et al*., 2012), with 5 of these 11 external to the networks of interest (spanning language, visual, auditory, cerebellar and somato-motor networks), while the remaining 9 components appeared to be artefactual or noise components. The remaining 6 ICA components were identified as sub-networks of CEN, DMN and SN and were isolated for further analyses. These include the

- Left CEN comprised of left dlPFC and parietal structures
- Bilateral CEN comprised of bilateral dlPFC and parietal structures,
- Dorsal DMN comprised of posterior cingulate cortex (PCC) and MPFC nodes,
- Ventral DMN comprised of retrosplenial cortex and medial temporal lobe structures,
- Anterior SN comprised of anterior insula and dorsal ACC nodes, and
- Posterior SN comprised of posterior insula.

The spatial patterns of these components are shown in Figure 5.

**Figure 5:**
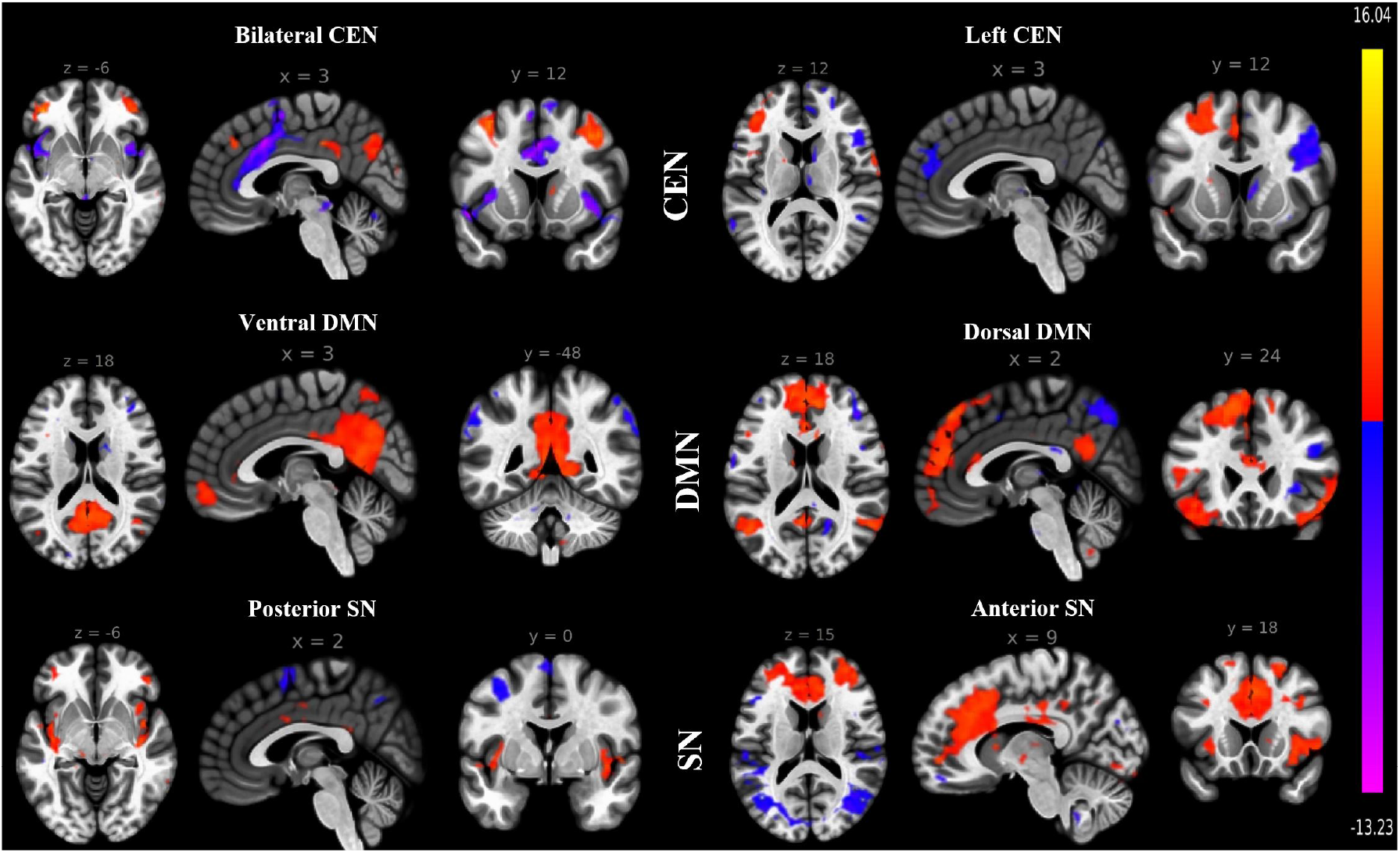
Spatial maps of ICA components with significant overlap with CEN, DMN and SN. Two ICA components were identified to have significant overlap with each one of the three networks.

### 3.2 Temporal characterization of tri-network activity

#### 3.2.1 ICA Networks - Timecourse Analysis

The ICA timecourses for the CEN, DMN and SN networks are shown in Figures 6a and 6b, averaged over all ABM and WM trials respectively.

**Figure 6:**
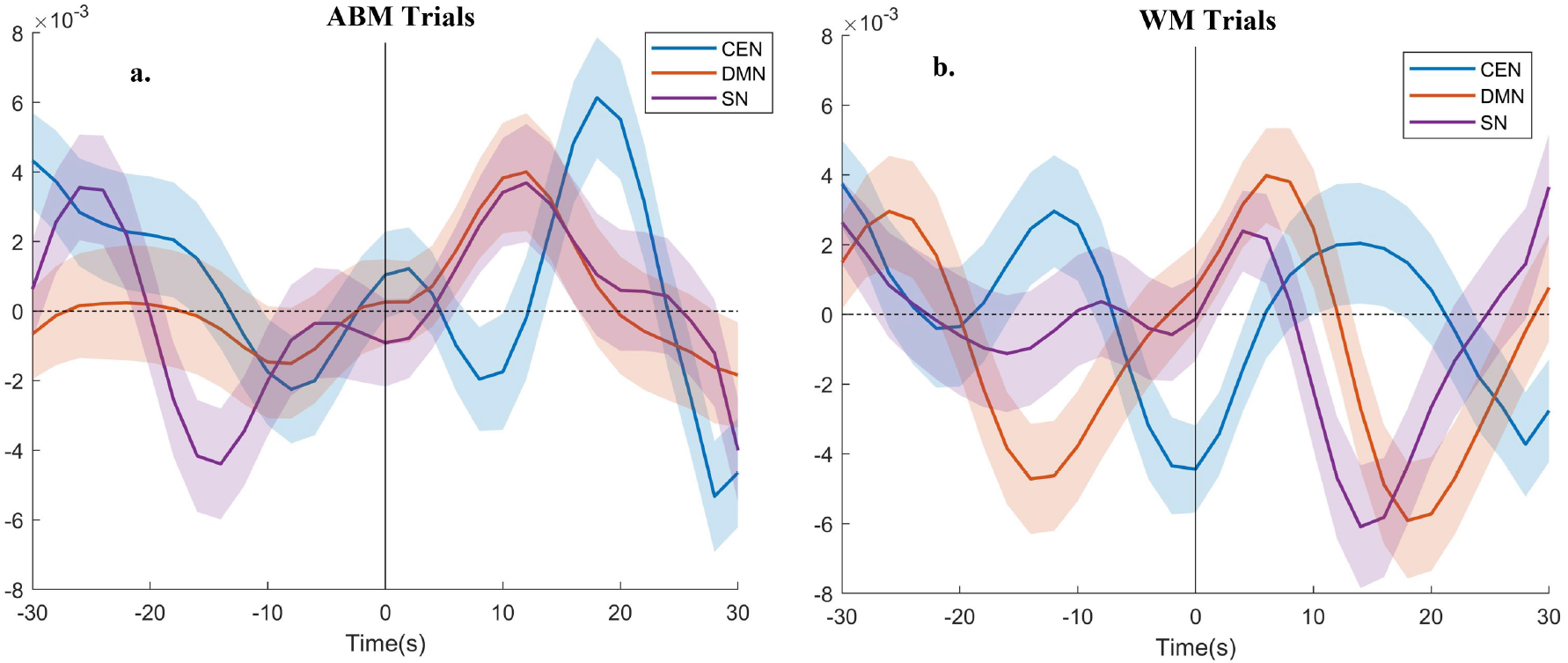
Activity timecourses of the ICA networks corresponding to the CEN (blue), DMN (orange) and SN (purple), averaged over all a. ABM trials, and b. WM trials. Time is shown with respect to the task onset (time = 0s).

During the ABM trials (Figure 6a), as predicted, the SN was observed to coactivate with the DMN, jointly increasing after t=0, and peaking at 12s post-task onset. At the same time, the CEN activity decreased through the first half of the ABM trials, with its lowest point coinciding with the DMN/SN activity peak. Surprisingly, after the point of peak DMN and minimum CEN activity at 12 s, DMN activity decreased while CEN activity increased, reaching its peak at 18s, suggesting there may be greater demands on executive functions during the latter half of the trial. This peak in CEN activity, however, was not accompanied by a peak in SN activity.

During WM trials, unexpectedly the SN did not seem to coactivate with the CEN. Instead, the activity of all three networks increased after t=0s, reaching their maximum levels at different times. The SN was the first to reach its peak at 5s, followed by the DMN at 8s, ending with the CEN peak at 13s.

The observed pattern of SN network activity was unexpected in both ABM and WM trials. We predicted that the SN would control the selection between the CEN and DMN according to task demands and therefore expected SN activity to rise at the start of each block, followed by activation of either the CEN or DMN in a task dependent manner. Additionally, we expected the DMN to be activated during ABM trials but not WM trials and the CEN to be activated during WM trials but not ABM trials, however, as noted above, there was cross-network activation in both types of trials. We postulated that this pattern of tri-network activity could be due to differential activity within its constituent sub-networks and we therefore investigated the anterior vs posterior sub-networks of the SN, alongside the CEN and DMN sub-networks identified by the group-wide ICA analysis, described in section 3.1.2. The activation timecourses of these SN, CEN and DMN subnetworks are shown in Figure 7.

**Figure 7:**
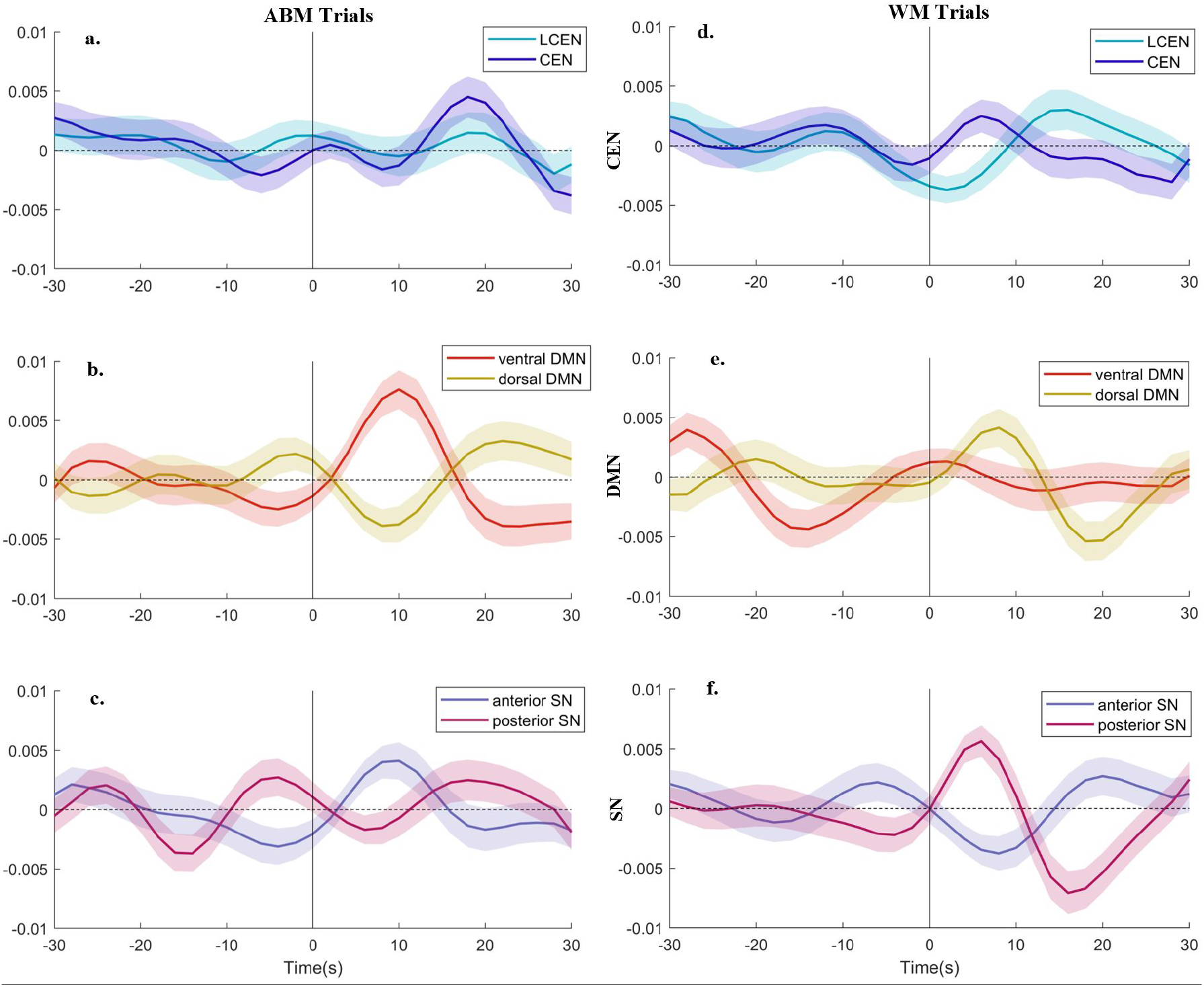
Activity timecourses of the ICA sub-networks corresponding to the CEN (light/dark blue), DMN (red/yellow) and SN (purple/violet), averaged over all ABM trials (panels a, b and c, respectively) and WM trials (panels d, e and f, respectively). Time is shown with respect to the task onset (time = 0s).

The correlated increase in DMN and SN activity during ABM trials observed in Figure 6a was found to be due to the correlated increase in activity of the ventral DMN and anterior SN sub-networks (seen in panels b and c respectively), whereas the later increase in CEN activity in Figure 6a was due to an increase in activity of the bilateral CEN network (panel a). In contrast, during the WM trials, the posterior SN (panel f) and the dorsal DMN activity (panel e) seemed to underlie the early SN and DMN peaks, while the LCEN subnetwork (panel d) contributed to the later CEN peak at 13s observed in Figure 6b. An early peak in bilateral CEN activity was also observed, coinciding with the increase in dorsal DMN and posterior SN activity in panels d, e and f of Figure 7, respectively.

In sum, two key patterns emerged in these sub-network analyses. Firstly, the anterior SN subnetwork showed co-activation with the task-relevant network (ventral DMN during ABM trials and LCEN during WM trials). In contrast, the posterior SN subnetwork was anti-correlated with the anterior SN, and instead consistently co-activated with the bilateral CEN and dorsal DMN subnetworks during both ABM and WM trials. Secondly, the LCEN co-activated with the bilateral CEN during the ABM trials, however, the LCEN showed a distinctly different activation pattern during WM trials, increasing in activity during the latter half of the trials (Figure 7d).

The network dynamics of these sub-networks were further studied using multivariate Granger causality (MVGC) analyses and are discussed later in section 3.3.1.

While the above analyses are based on the data-driven (ICA) identification of the networks and sub-networks, we also examined the tri-network activation in terms of ROI analyses in the next section.

#### 3.2.2 ROI-to-ROI Timecourse Analysis

In addition to ICA based tri-network definitions, we investigated the tri-network activation using two different sets of ROIs to describe the CEN, DMN and SN. The first network configuration utilized an abridged set of ROIs widely used in the literature to study the tri-network model, while the second network configuration extended the set of ROIs defining the DMN and SN to include medial temporal lobe (MTL) and posterior insula (PI) nodes. The results from these ROI analyses are discussed in this section.

The average BOLD activity timecourses of each of the three major networks, gathered by summing together the activity of the corresponding nodes, is shown in Figures 8a and 8b, averaged across all ABM and WM task trials respectively.

**Figure 8:**
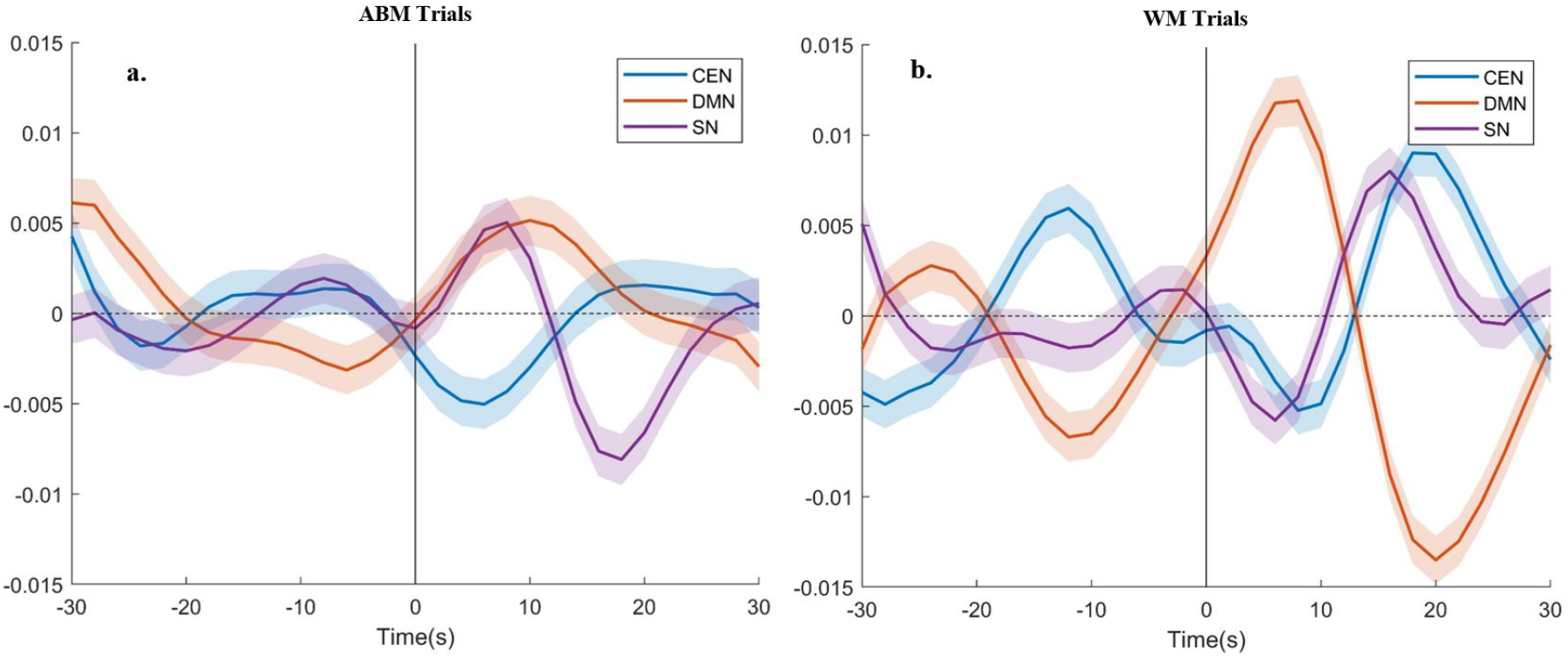
BOLD activity timecourses of the CEN (blue), DMN (purple) and SN (orange), computed from activity of the corresponding nodes, averaged over all a. ABM trials, and b. WM trials. Time is shown with respect to the task onset (time = 0s). BOLD activity is shown in units of percent signal change from mean activity (represented by the dotted line).

During the ABM trials, consistent with our findings from the ICA-based sub-network analyses, SN activity was the first to increase, reaching its peak at 8 seconds, followed by an increase in DMN activity which peaked at 11 seconds. As expected, CEN activity decreased by the same amount during the same time frame, and increased only after the DMN activity had returned to baseline around 18 seconds post-task onset. This change in CEN activity was accompanied by a decrease in SN activity at 18 seconds post-task onset. During the WM trials, the DMN activity increased rapidly, peaking around 6 seconds, while the SN and CEN activity decreased during the same time frame. This was followed by an increase in SN and CEN activity, peaking at 17 seconds and 19 seconds respectively. This was accompanied by a large decrease in DMN activity, reaching its lowest activity level at 20 seconds post-task onset. Across both ABM and WM trials, the SN was found to co-activate with the activity of the network predominantly expected to activate, i.e. DMN during the ABM trials and CEN during the WM trials. Furthermore, SN activity was observed to lead the activity of the activated network, consistent with the activation pattern expected for a network gating the onset of another network.

The activity of each network was further broken down into the activity of its constituent nodes for more detailed analysis, as shown in Figure 9. During the ABM trials, there was an increase in vmPFC activity at 5 seconds post-task onset (panel b), followed by dACC, right AI and left AI activity around 8 seconds post-task onset (panel c), after which PCC activity increased to its maximum value around 10 seconds post-task onset (panel b). The amPFC activity followed an opposite trend to that of PCC (panel b). During the latter half of the trial, right and left dlPFC activity were found to increase around 18 seconds post-stimulus onset (panel a). During the WM trials, the activity of SN and CEN nodes was found to be biphasic. First, the dACC activity increased at 14 seconds post-stimulus onset (panel f), immediately followed by an increase in right PPC activity (panel d). In the second phase, the right and left AI activity increased to their maximum values at 16 seconds and 20 seconds post-task onset respectively (panel f). This was followed by an increase in left PPC, left dlPFC and finally right dlPFC, peaking at 18, 20 and 22 seconds respectively (panel d). Activity of the DMN nodes during the WM trials was dominated by an early increase in PCC activity around 7 seconds (panel e), which was accompanied by a decrease in right and left AI node activity (panel f).

**Figure 9:**
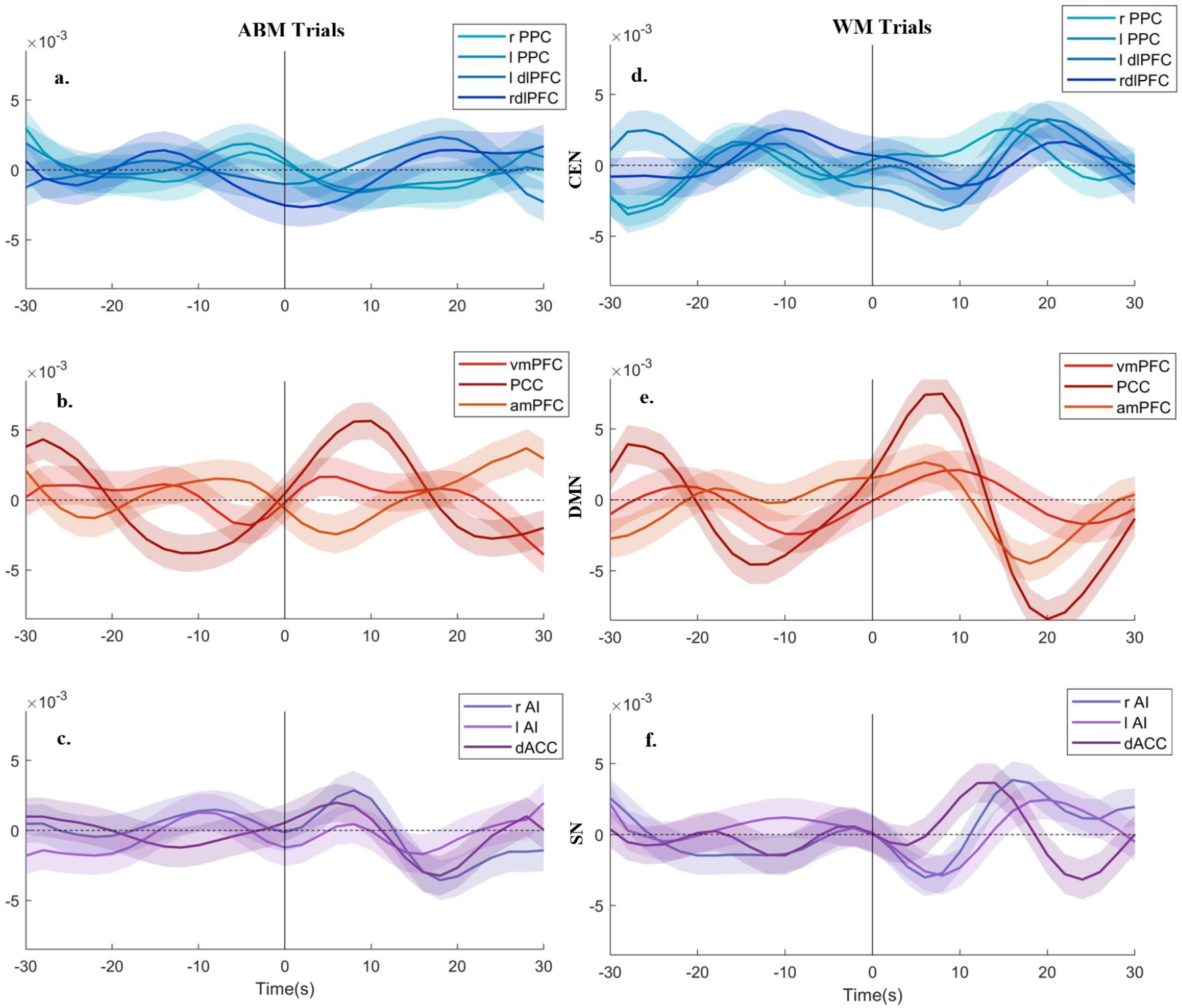
Activity timecourses of the ROIs corresponding to the CEN (light/dark blue), DMN (red/yellow) and SN (purple/violet) networks, using the abridged node definitions, averaged over all ABM trials (panels a, b and c, respectively) and WM trials (panels d, e and f, respectively). Time is shown with respect to the task onset (time = 0s).

Taken together, the results described above and illustrate in Figures 6 and 8 suggest that different nodes dominated the activity of each network during the two different tasks. The observed increase in DMN activity during the ABM task might have been due to an increase in vmPFC and PCC activity driven by rAI, lAI and dACC activity, whereas, the early increase in DMN activity during the WM trials was primarily due to a corresponding increase in PCC and amPFC activity. This increase in PCC activity during WM trials was accompanied by a decrease in SN node activity indicating that this activation might be negatively correlated to the SN. Furthermore, the similar patterns of PCC activity observed during both ABM and WM trials might indicate some common component processes required to adequately complete both tasks. The late increase in CEN activity during the ABM task was dominated by an increase in right and left dlPFC activity, and was also accompanied by a decrease in SN node activity. However, the increase in net CEN activity during the WM task was found to be due to an underlying increase in right PPC activity and that of the dACC node of the SN, followed by an increase in left PPC and bilateral dlPFC activity, shortly preceded by an increase in bilateral AI activity. Consequently, this CEN activation might have been driven by the SN. To better investigate the causal patterns underlying the observed network dynamics, multivariate Granger causality (MVGC) analysis was performed using the described network nodes, and its results are discussed in section 3.3.2.

While the above described results rely on a small subset of nodes used to describe the CEN, DMN and SN in prior studies investigating the tri-network model (abridged ROIs), they omit a key set of medial temporal lobe (MTL) nodes implicated in DMN-linked memory tasks, in addition to the posterior insula (PI), which is thought to participate in the bottom-up salience detection functions of the SN. To investigate the impact of including these key nodes within the network definitions on tri-network activity, we added right and left hippocampus (r/l HC), right and left parahippocampus (r/l pHC) and the retrosplenial cortex (RSC) to the set of DMN nodes. Additionally, we also included the right and left posterior insula (r/l PI) in the set of SN nodes. The impact of including the extended set of nodes on tri-network activity is discussed below.

During the ABM trials, the addition of the MTL nodes in the DMN network definition introduced a second peak in DMN activity in the latter half of the trials around 18s posttask onset (Figure 10a). This peak coincided with the peak in CEN activity during the ABM trials and was not accompanied by a corresponding SN peak. This second peak could be attributed to the increase in activity within the MTL nodes, as seen in Figure 11b. Interestingly, this late DMN peak was also accompanied by an increase in PI activity (panel c), although this increase was not sufficient to increase the overall SN activity seen in Figure 10a. In addition to this late peak in DMN activity, the MTL nodes also showed an increase in activity during the first 5-6 seconds of the trial, co-activating with the increase in PCC activity during the same time period (panel b).

**Figure 10:**
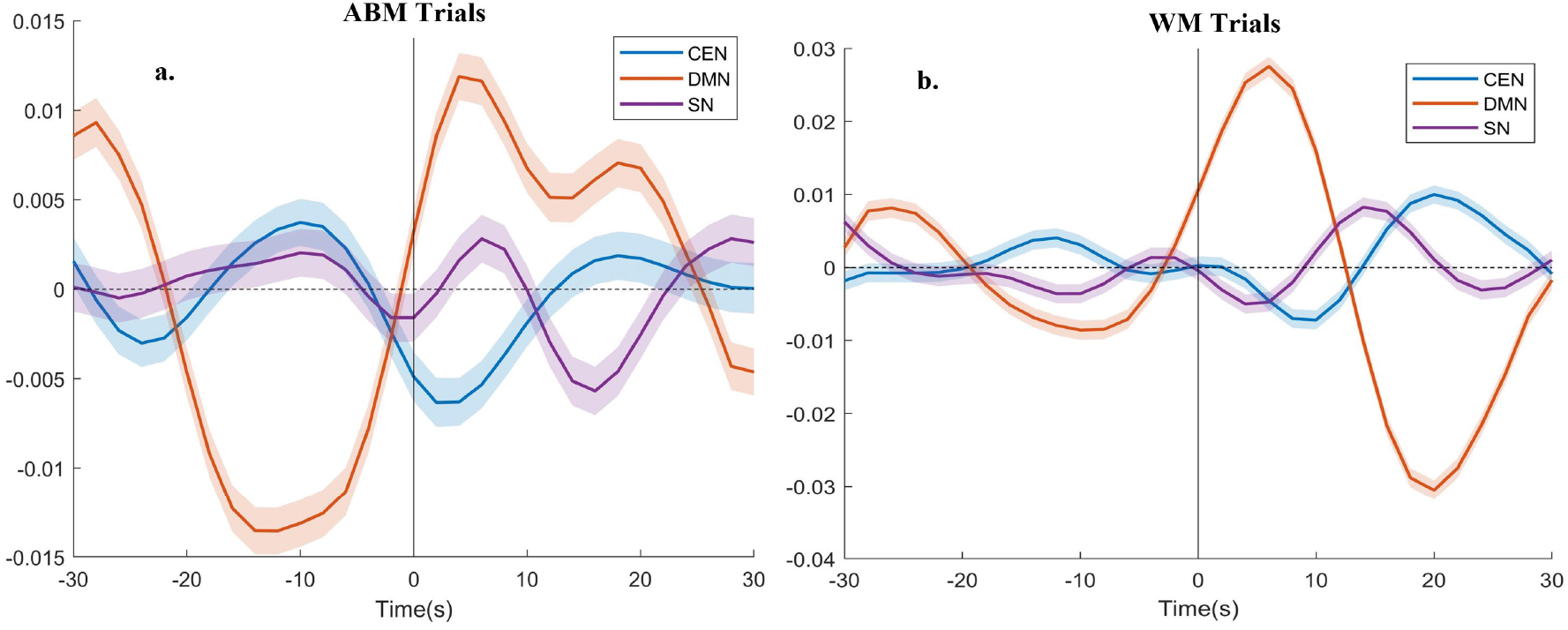
BOLD activity timecourses of the CEN (blue), DMN (purple) and SN (orange), computed from activity of the corresponding nodes using the extended set of ROIs, averaged over all a. ABM trials, and b. WM trials. Time is shown with respect to the task onset (time = 0s). BOLD activity is shown in units of percent signal change from mean activity (represented by the dotted line).

**Figure 11:**
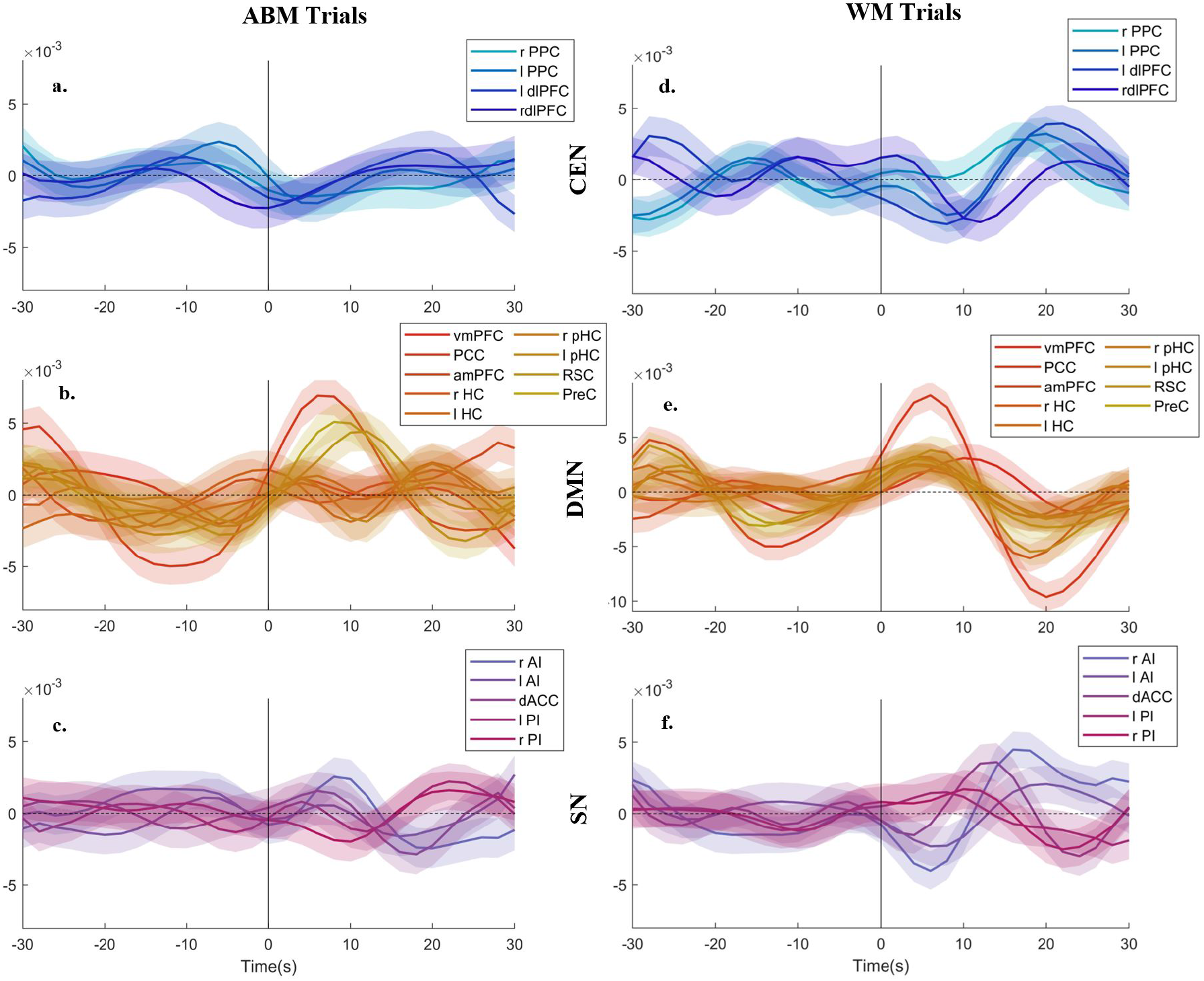
Activity timecourses of the ROIs corresponding to the CEN (light/dark blue), DMN (red/yellow) and SN (purple/violet) networks, using the extended node definitions, averaged over all ABM trials (panels a, b and c, respectively) and WM trials (panels d, e and f, respectively). Time is shown with respect to the task onset (time = 0s).

This early increase in MTL node activity was also observed during the WM trials, however, in contrast to the ABM trials, there was no increase in MTL activity in the latter half of the trials (panel e). The PI activity was observed to be elevated during the early increase in MTL node activity, followed by a decrease in activity during the latter half of the WM trials dominated by bilateral AI activity (panel f).

In conclusion, the findings including the extended set of DMN and SN nodes indicate the involvement of key memory-linked MTL nodes in the cognitive processes required during the ABM and WM trials. A common pattern of increased MTL activity was observed during the first half of both ABM and WM trials, that correlated with a similar increase in PCC and PreC activity, potentially indicating that some of the common component processes between the two tasks relied on key MTL nodes and could be related to memory processing.

### 3.3 Causality analysis of tri-network activity

To appropriately understand the causal influence of the above described sub-networks and ROIs, we conducted multivariate Granger causality analyses and describe them below.

#### 3.3.1 ICA Networks - Multivariate Granger Causality (MVGC) Analysis

MVGC analyses revealed that the optimal model order for estimating the VAR was 3-4 time points using an AIC criterion, implying that lagging signals most likely had a lag of 6 to 8 seconds (model order x TR). Furthermore, the MVGC connectivity patterns generally supported the observations made in the previous sections.

##### ABM Trials

During the onset of the ABM Trials, the strength of the pairwise-conditional causal outflow from the anterior SN sequentially increased to the left CEN, bilateral CEN, the ventral DMN and then the dorsal DMN. The anterior SN to ventral DMN direction (Figure 12) causality was at its peak around the same time as the peak in ventral DMN activity (Figure 7b) post-task onset, also seen in the sub-network graphs from time t = 1s through t = 7s in Figure 12. This observed increase in causality accompanying the correlated increase in anterior SN and ventral DMN activity could be indicative of the anterior SN recruiting the ventral DMN sub-network post-ABM task onset.

**Figure 12:**
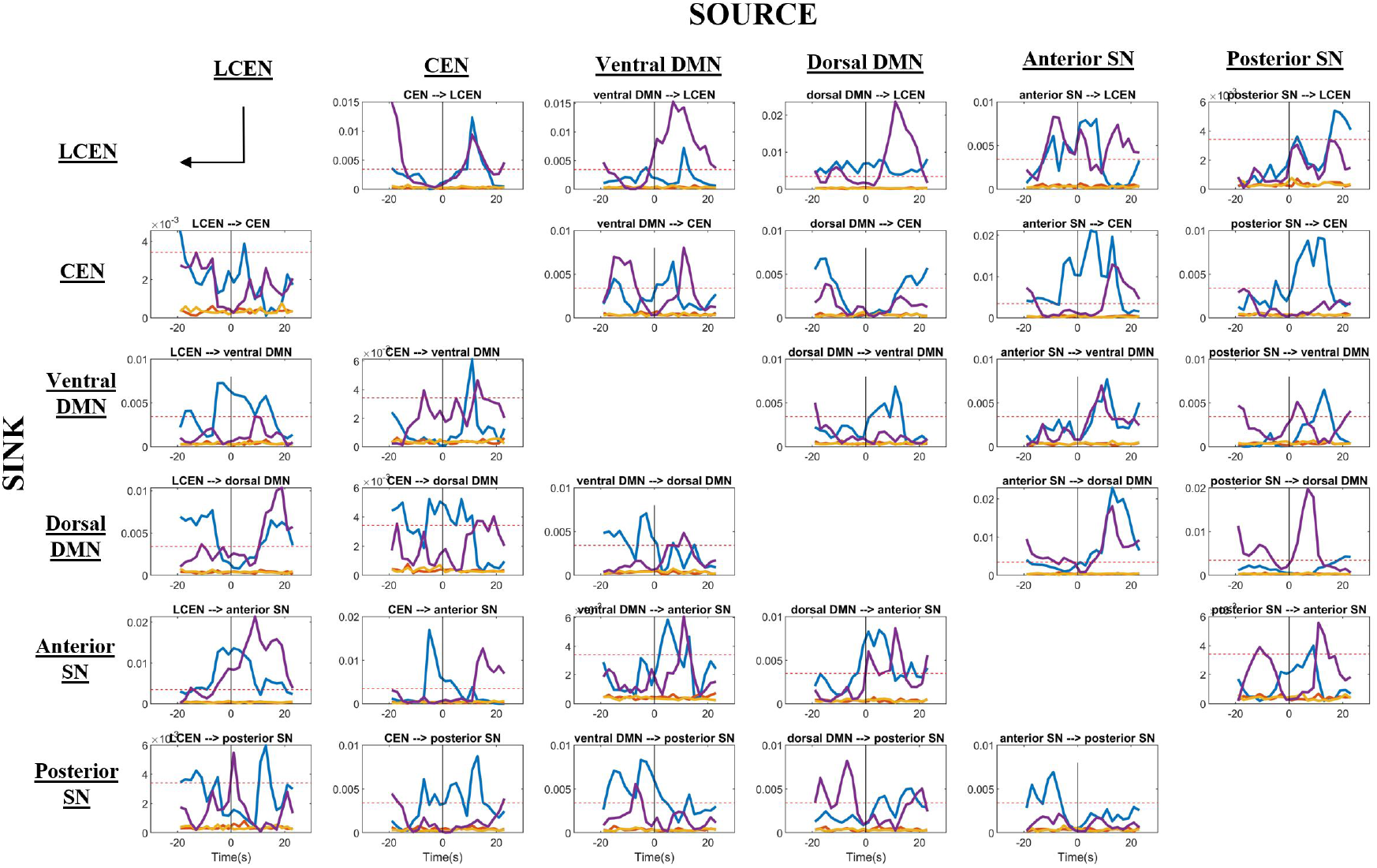
MVGC values between the six sub-networks, as they vary across the ABM and WM trials, shown in blue and purple respectively. Each MVGC value is estimated using sliding window of width 14 seconds and step 2 seconds. Null MVGC values are also shown (orange) along with the threshold for FDR-corrected significance at p<0.05 (dotted red line). Each column is a source node, while each row is a sink node, with each row-column intersection representing the connection going from the corresponding source node to the sink node.

Net causal inflow to the ventral DMN increased during the first few seconds (t=1s to t=15s) post-task onset, primarily due to an increase in anterior SN to ventral DMN causality. It was also supplemented by an early increase in input from the posterior SN (t=1s to 3s), followed by an increase in bilateral CEN input (t=3s to 7s). The direction of net causal flow to the ventral DMN switched from primarily causal inflow to primarily causal outflow around t=7s, and lasted until t= 17s. This pattern of causal flow might represent the recruitment of the ventral DMN sub-network by the anterior SN during the first few seconds of the ABM trials, followed by DMN-generated recruitment of other regions during AM memory retrieval and elaboration.

The peak in anterior SN to ventral DMN causality was preceded by a biphasic peak in anterior SN to bilateral CEN causality, followed by a peak in anterior SN to dorsal DMN causality. However, the activity of these sub-networks were anti-correlated with that of the anterior SN (Figure 7), potentially indicative of the anterior SN’s role in suppressing dorsal DMN and bilateral CEN activity when recruiting the ventral DMN during the ABM trials.

The dorsal DMN sub-network was found to have a mix of incoming and outgoing connections to other sub-networks shortly after ABM-task onset, followed by predominantly incoming connections from the left CEN, anterior SN and posterior SN during the latter half of the ABM trials (t=17s onwards), as seen in the corresponding panels in Figure 13. These causal patterns for the dorsal DMN are distinctly different from that of the ventral DMN, indicating different ABM-linked activity in the nodes comprising these sub-networks.

**Figure 13:**
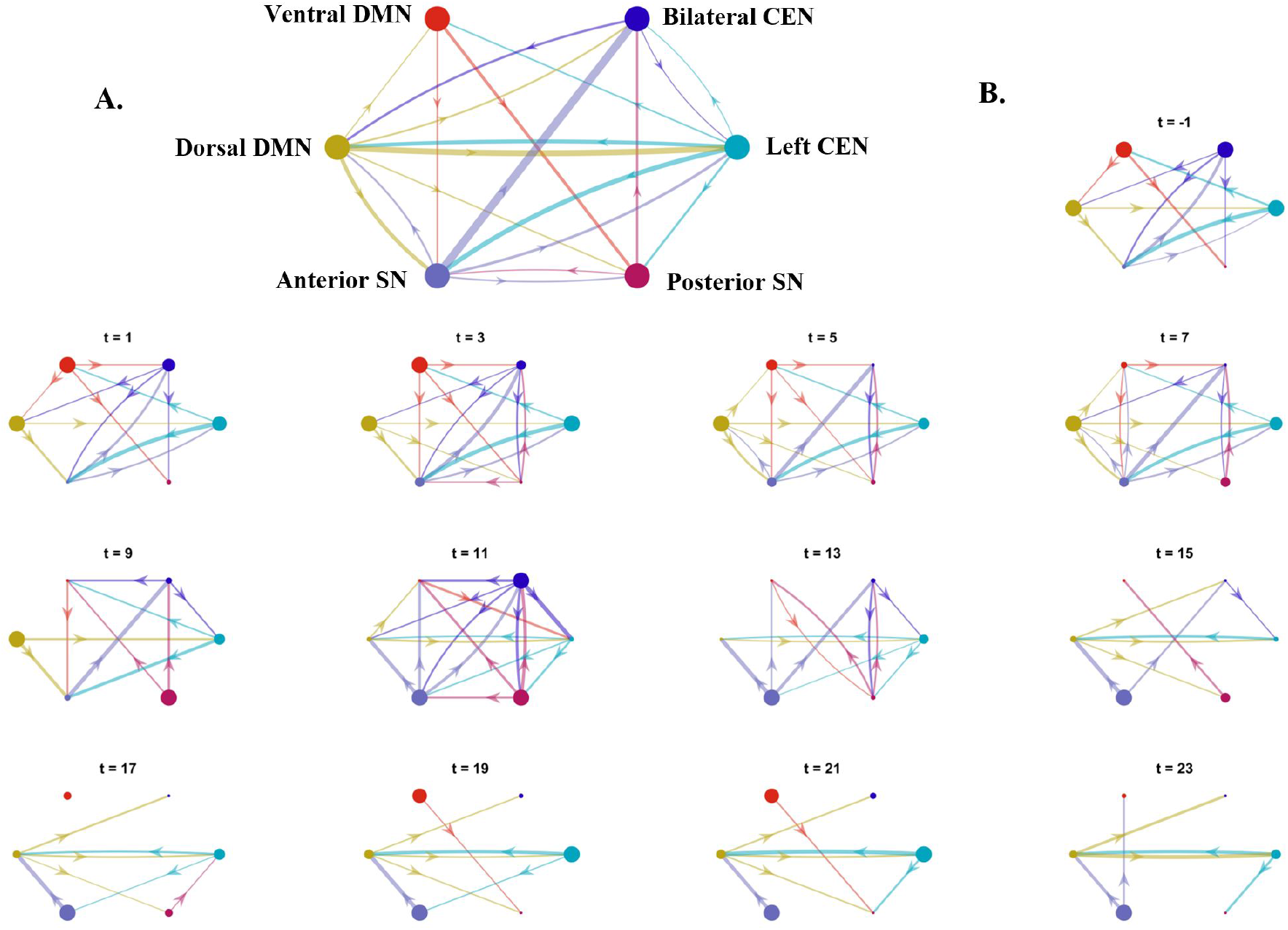
MVGC connections between the six sub-networks, shown for all ABM trials. Connections passing FDR correction at p < 0.05 level are shown. A. The MVGC connections estimated using the entire trial duration; B. The MVGC connections estimated using a sliding window of width 14 seconds and step 2 seconds. The time label for each graph corresponds to the center of each window. The sizes of each node is scaled according to the MVGC weighted out-degree/in-degree. The thickness of each edge is scaled according to the MVGC value representing the edge.

**Figure 14:**
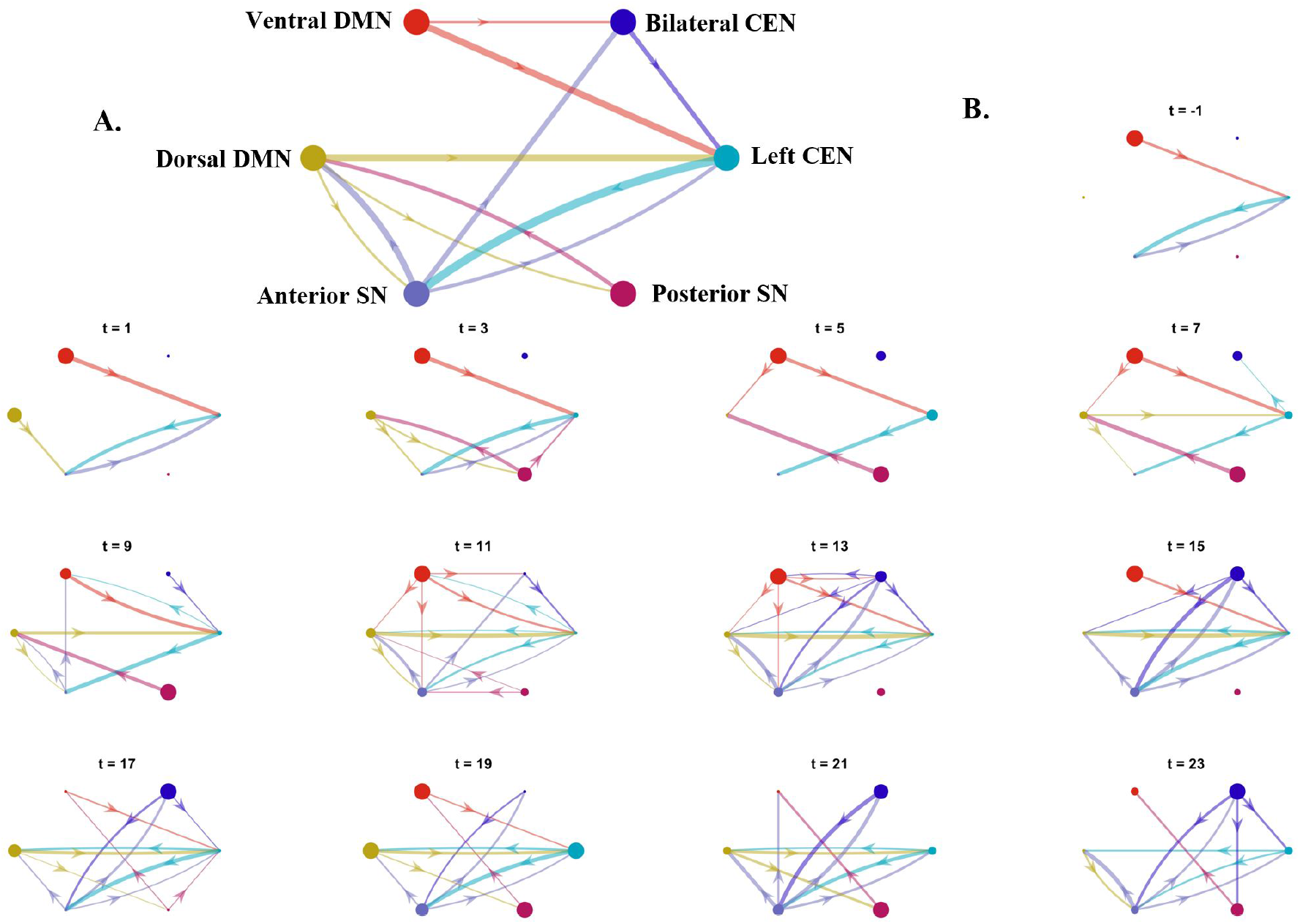
MVGC connections between the six sub-networks, shown for all WM trials. Connections passing FDR correction at p < 0.05 level are shown. A. The MVGC connections estimated using the entire trial duration; B. The MVGC connections estimated using a sliding window of width 14 seconds and step 2 seconds. The time label for each graph corresponds to the center of each window. The sizes of each node is scaled according to the MVGC weighted out-degree/in-degree. The thickness of each edge is scaled according to the MVGC value representing the edge.

##### WM Trials

After WM task onset, the pairwise conditional causality from anterior SN, ventral DMN and posterior SN to left CEN increased. The causality from left CEN to bilateral CEN also increased along with a concomitant increase in bilateral CEN activity (Figure 7d). Furthermore, the causality from posterior SN to dorsal DMN also increased along with an increase in dorsal DMN activity. These changes in pairwise causality contributed to the co-activation of bilateral CEN, dorsal DMN and the posterior SN during the first 10 seconds of WM trials (Figure 7 d, e and f). This pattern is also observed in the latter half of the ABM trials, potentially indicative of CEN-mediated working memory maintenance processes common to both ABM and WM trials.

The pairwise-conditional causality from ventral DMN to left CEN also increased close to the task onset, continuing to increase until 10 s. Accompanied by an increase in causal inflow from dorsal DMN and bilateral CEN, this peak in causal input to the left CEN coincided with an increase in left CEN activity (Figure 7d), which persisted through the rest of the trial. This causal pattern is indicative of left CEN activation, potentially for the maintenance of working memory.

In sum the above results suggest the anterior SN sub-network influenced the key task-relevant networks, while the posterior SN sub-network consistently interacted with working memory-linked sub-networks common to both ABM and WM tasks. The observed causal patterns also emphasise the different functional roles of the bilateral CEN and left CEN sub-networks, potentially involved in initializing working memory processes and the maintenance of working memory, respectively.

#### 3.3.2 ROI-to-ROI - Multivariate Granger Causality (MVGC) Analysis

In addition to exploring the causality of data-driven ICA sub-networks, we conducted MVGC analysis on the ROI-to-ROI data to understand causal interactions between an abridged set of widely described nodes of tri-network model, as well as an extended set of nodes representing the tri-network model.

The AIC criterion analysis found 4 time points to be the optimal lag for the VAR model used in the MVGC analysis. This corresponded to an average time lag of 8 seconds for the pairwise conditional Granger causality between two nodes. The timecourses of the MVGC estimates between each pair of nodes for the ABM and WM trials are shown in Figure 15. The connectivity between this abridged set of tri-network nodes is further visualized from t = −1 seconds to t = 23 seconds post-task onset for the ABM and WM trials in Figures 16 and 17 respectively. This is contrasted with the temporal changes in connectivity between the extended set of tri-network nodes, shown in Figures 18 and 19 for the ABM and WM trials respectively. The size of each node in these graphs corresponds to their net causal outflow (outflow - inflow), which is shown separately for the ABM and WM trials in Figure 20. The hubness score of each node is also shown in Figure 21. Key observations from these results are discussed below.

**Figure 15:**
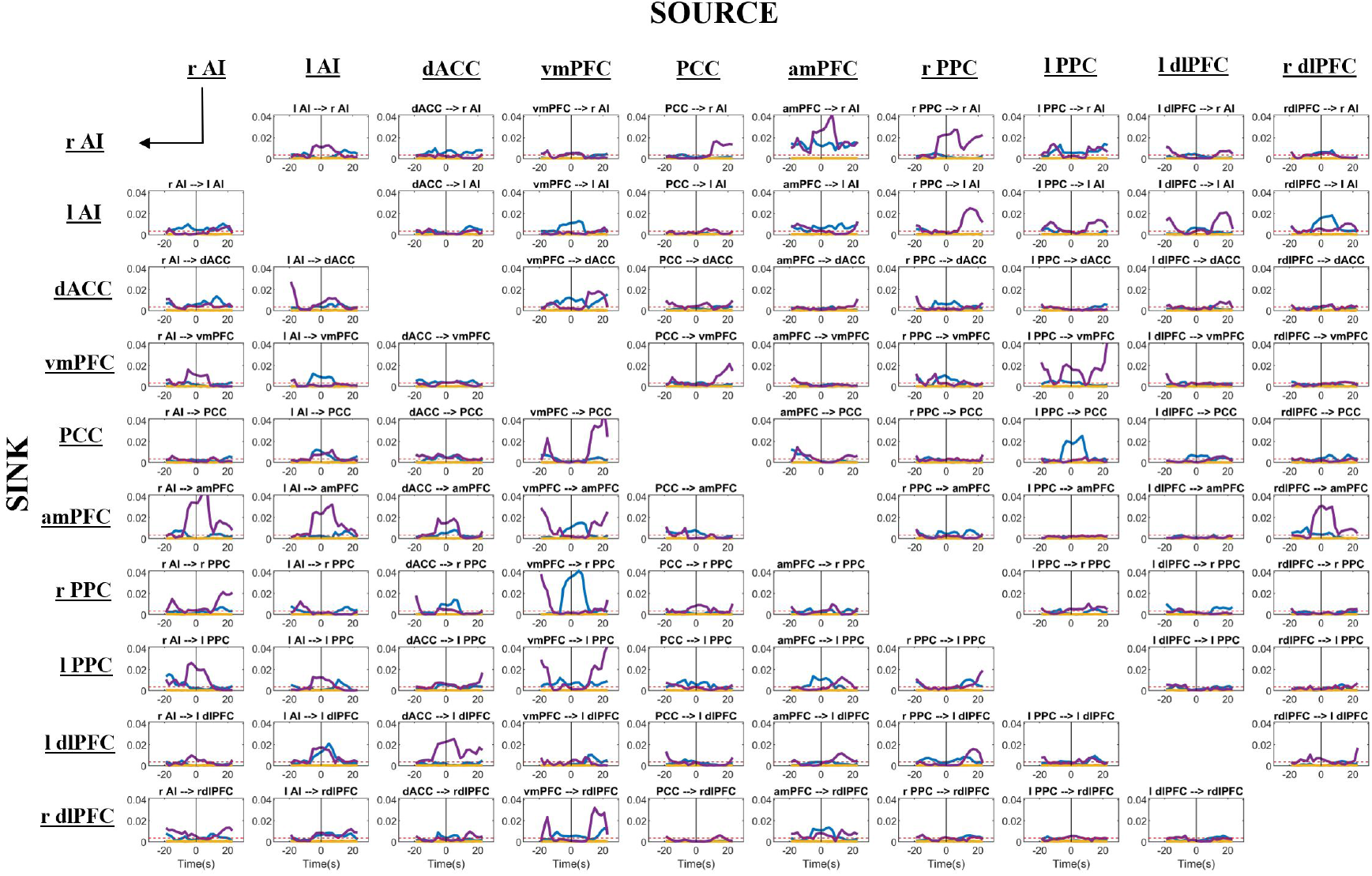
MVGC values between each pair of the network nodes, as they vary across the ABM and WM trials, shown in blue and purple respectively. Each MVGC value is estimated using a sliding window of width 14 seconds and step 2 seconds. Null MVGC values are also shown (orange) along with the threshold for FDR-corrected significance at p<0.05 (dotted red line). Each column is a source node, while each row is a sink node, with each row-column intersection representing the connection going from the corresponding source node to the sink node.

**Figure 16:**
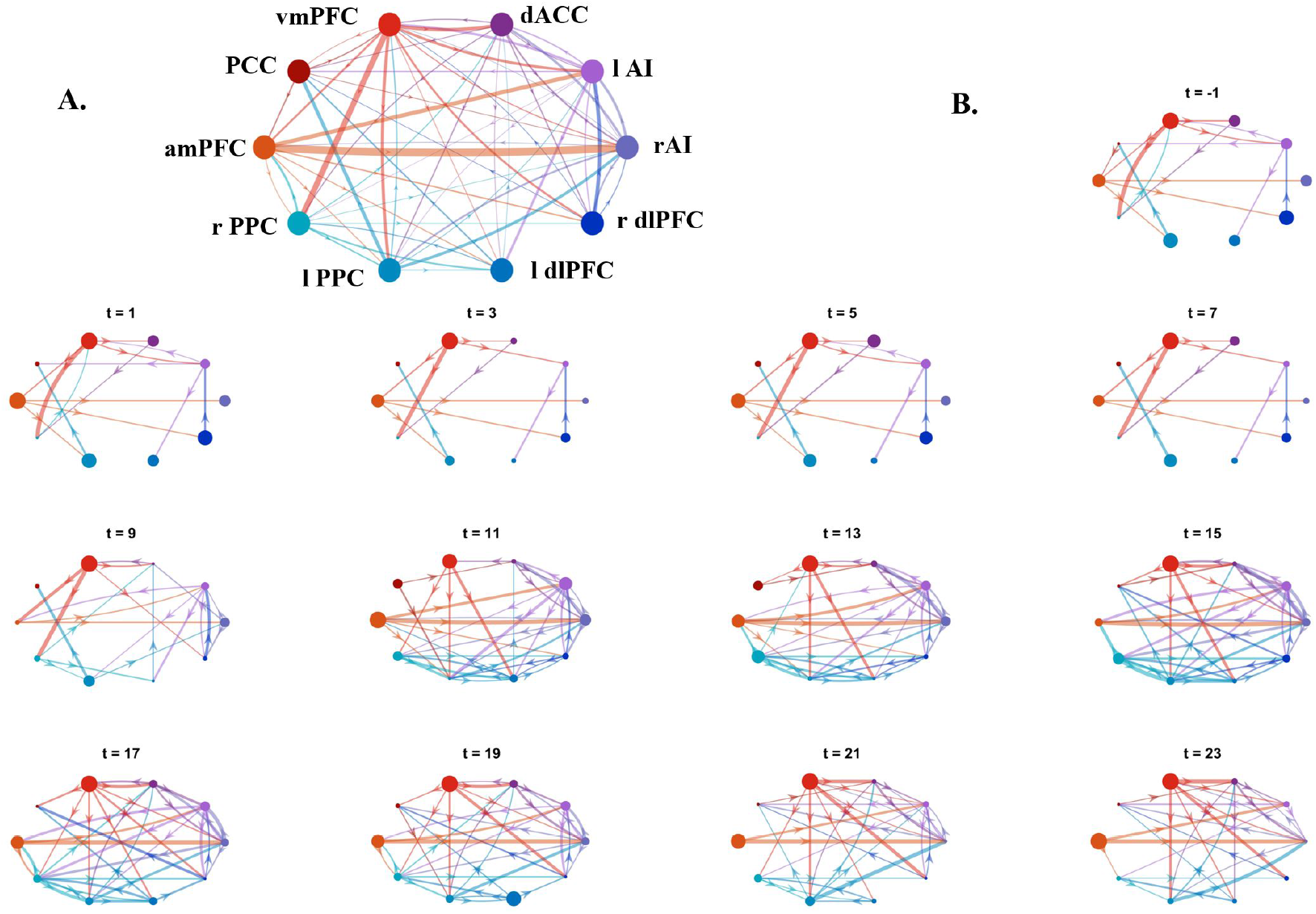
MVGC connections between the network nodes, shown for all ABM trials. Only connections passing FDR correction at p < 0.05 level are shown. A. The MVGC connections estimated using the entire trial duration; B. The MVGC connections were estimated using a sliding window of width 14 seconds and step 2 seconds. The time label for each graph corresponds to the center of each window. The size of each node is scaled according to the MVGC weighted out-degree/in-degree, whereas the thickness of each edge is scaled according to the magnitude of the corresponding MVGC value. Furthermore, the color of each edge corresponds to that of the source node and, along with the direction of the arrowhead, represents the edge direction.

**Figure 17:**
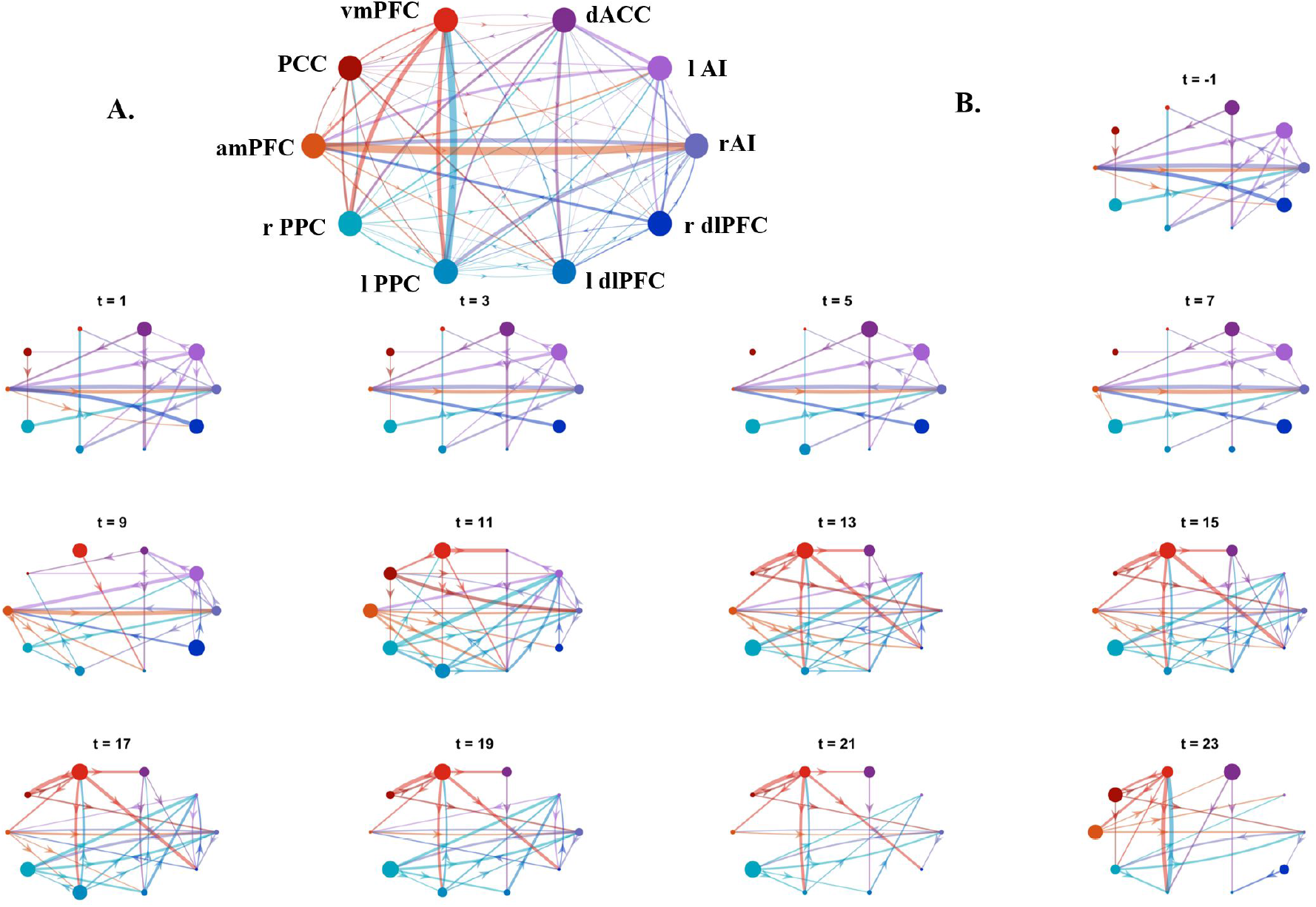
MVGC connections between the the network nodes, shown for all WM trials. Only connections passing FDR correction at p < 0.05 level are shown. A. The MVGC connections estimated using the entire trial duration; B. The MVGC connections were estimated using a sliding window of width 14 seconds and step 2 seconds. The time label for each graph corresponds to the center of each window. The sizes of each node is scaled according to the MVGC weighted out-degree/in-degree, whereas the thickness of each edge is scaled according to the magnitude of the corresponding MVGC value. Furthermore, the color of each edge corresponds to that of the source node and, along with the direction of the arrowhead, represents the edge direction.

**Figure 18:**
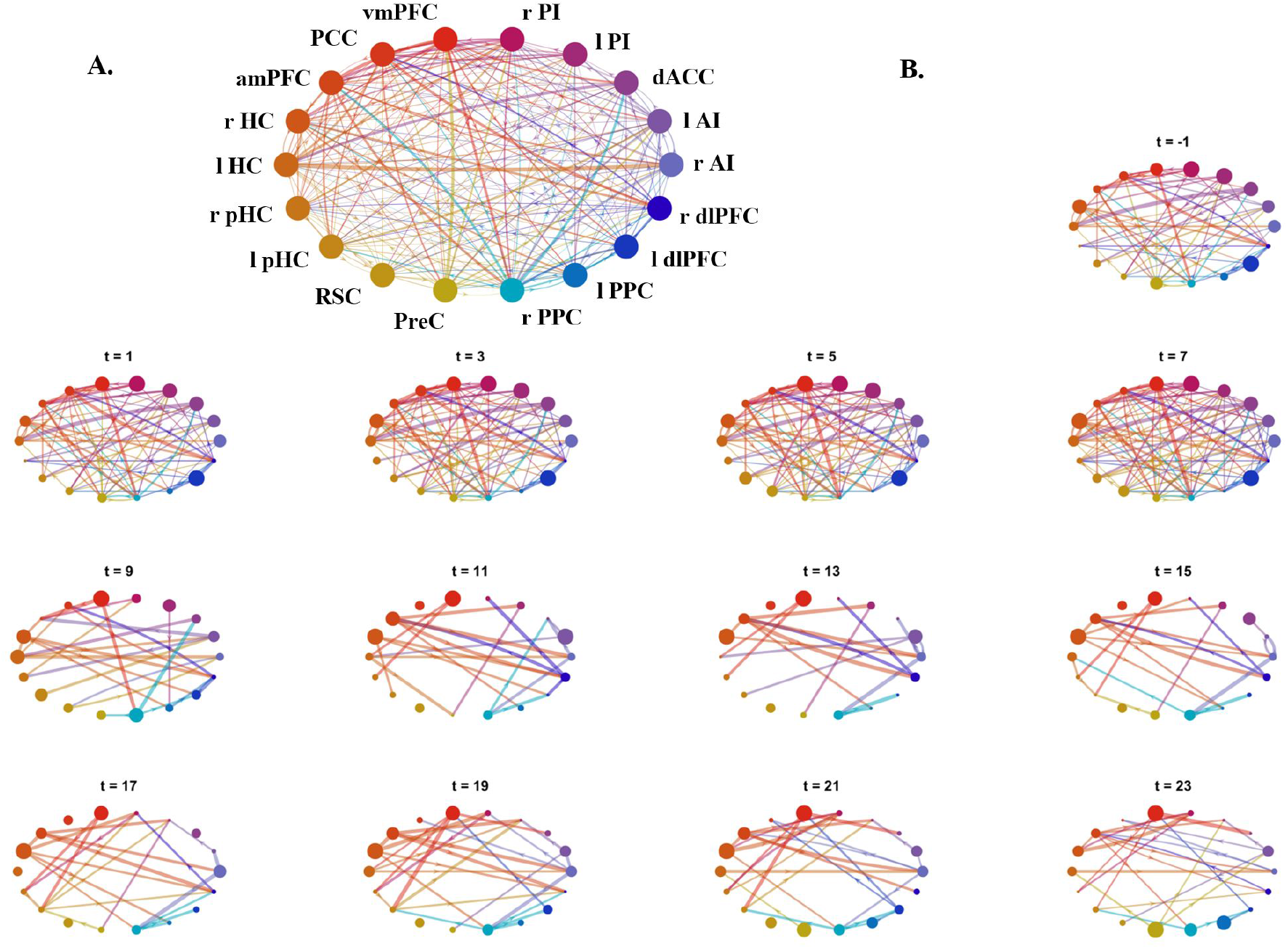
MVGC connections between the extended set of network nodes, shown for all ABM trials. Only connections passing FDR correction at p < 0.05 level are shown. A. The MVGC connections estimated using the entire trial duration; B. The MVGC connections were estimated using a sliding window of width 14 seconds and step 2 seconds. The time label for each graph corresponds to the center of each window. The sizes of each node is scaled according to the MVGC weighted out-degree/in-degree, whereas the thickness of each edge is scaled according to the magnitude of the corresponding MVGC value. Furthermore, the color of each edge corresponds to that of the source node and, along with the direction of the arrowhead, represents the edge direction.

**Figure 19:**
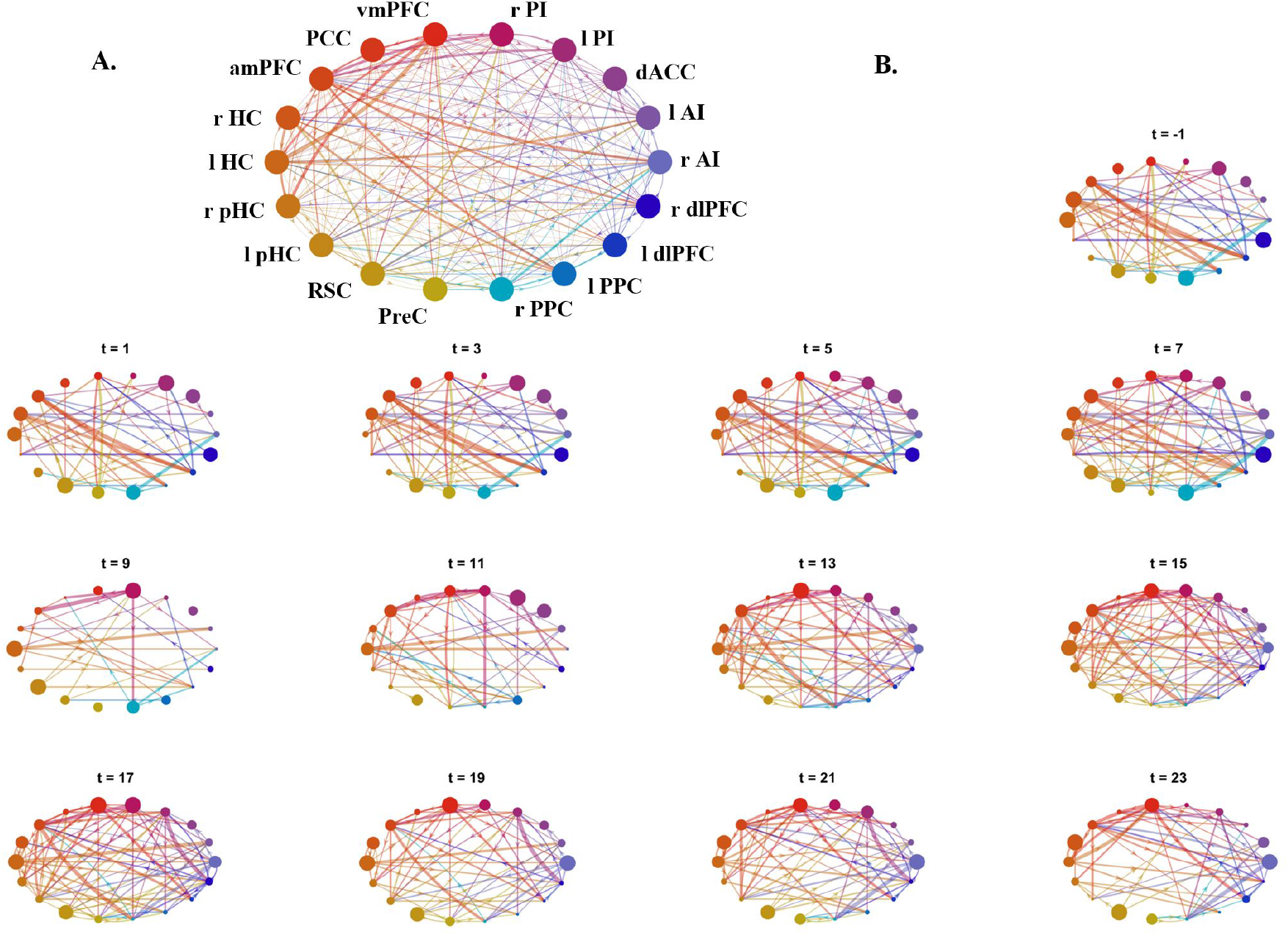
MVGC connections between the extended set of network nodes, shown for all WM trials. Only connections passing FDR correction at p < 0.05 level are shown. A. The MVGC connections estimated using the entire trial duration; B. The MVGC connections were estimated using a sliding window of width 14 seconds and step 2 seconds. The time label for each graph corresponds to the center of each window. The sizes of each node is scaled according to the MVGC weighted out-degree/in-degree, whereas the thickness of each edge is scaled according to the magnitude of the corresponding MVGC value. Furthermore, the color of each edge corresponds to that of the source node and, along with the direction of the arrowhead, represents the edge direction.

**Figure 20:**
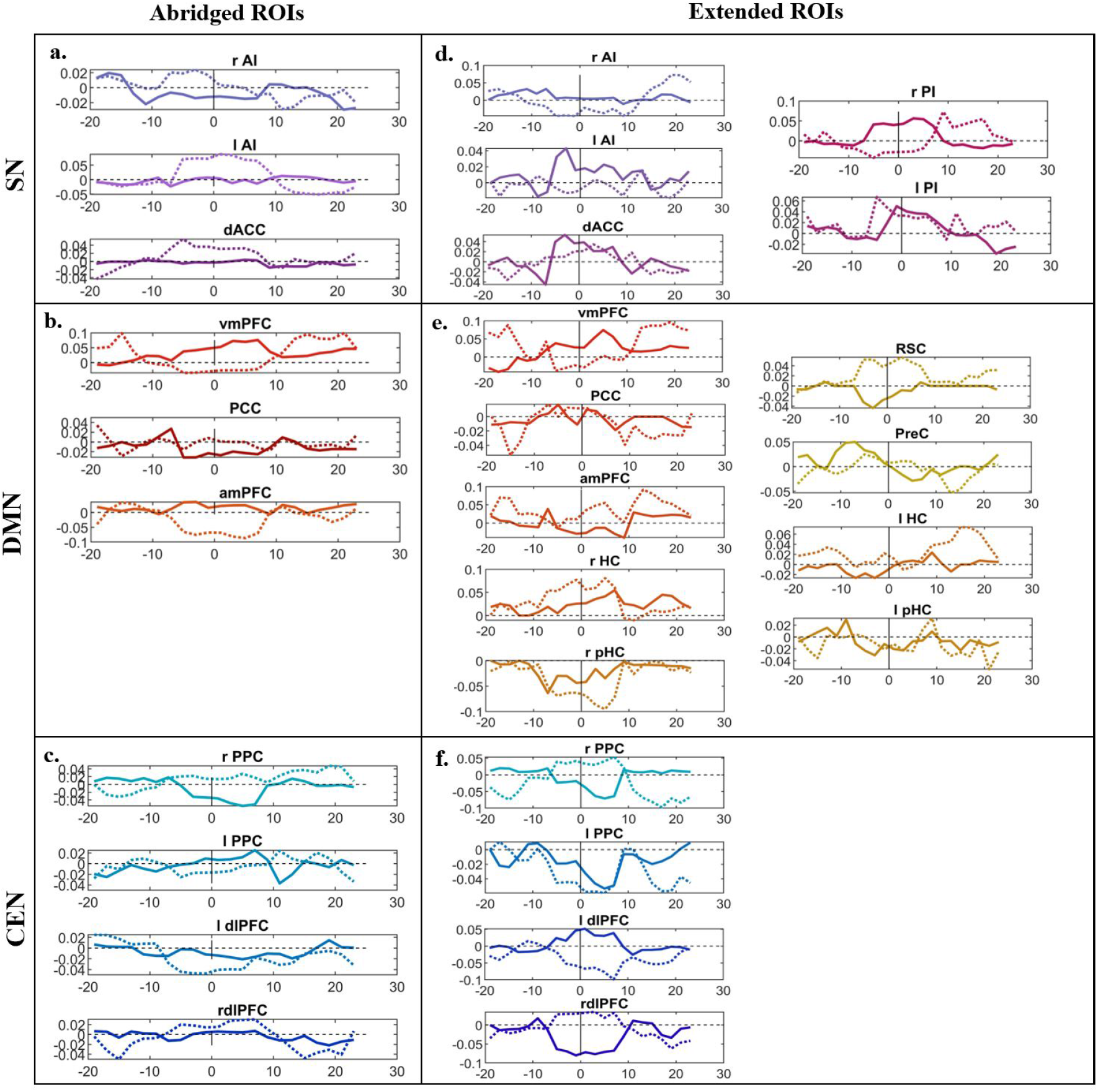
The net pairwise causal outflow from each network node (causal outflow - causal inflow) for the abridged set of a. SN, b. DMN and c. CEN nodes, alongside the causal outflow from the extended set of d. SN, e. DMN and f. CEN nodes. Each panel shows the time course of the causal outflow from a single network node, during the ABM (solid line) and WM trials (dashed line). A positive value corresponds to a new causal outflow and a negative value corresponds to a net causal inflow. Note the different range of causal outflow values for each network node. This allows for adequate visualization of the temporal patterns in causal outflow for all network nodes.

**Figure 21:**
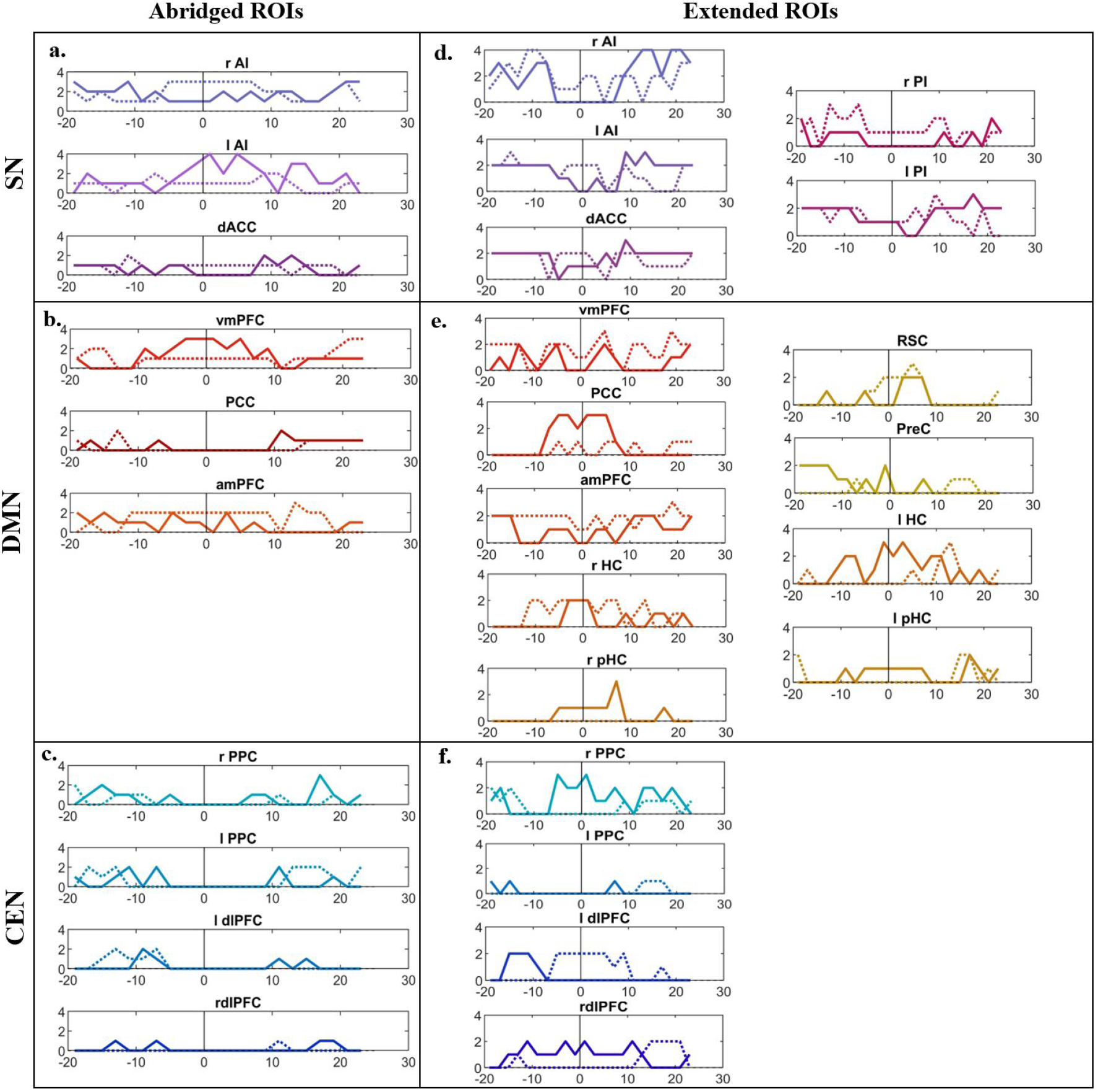
The hubness score for each network node for the abridged set of a. SN, b. DMN and c. CEN nodes, alongside the hubness score for the extended set of d. SN, e. DMN and f. CEN nodes. Each panel shows the time course of the hubness score from a single network node, during the ABM (solid line) and WM trials (dashed line). The hubness scores range from 0 to 4 and is scored according to the procedure described in section 2.3.4.

##### ABM Trials

The DMN node with maximal outgoing causal influence during the ABM trials was the vmPFC for the model using the abridged set of tri-network nodes, with outgoing connections to dACC and left AI of the SN during the first 8-10 seconds posttask onset (solid lines in Figure 20b). During this time period, a bi-directional connection between vmPFC and the right PPC of the CEN was also observed, with the vmPFC to right PPC direction being much stronger than the rPPC to vmPFC direction. Combined with the peak in vmPFC activity (Figure 9b) and the high hubness score (solid line in Figure 21b) during the same time period, these causal patterns might represent vmPFC-driven memory processes that also rely on posterior parietal cortex and left anterior insular nodes.

While the amPFC showed similar, albeit weaker causal influence on other nodes, the observed casual pattern of the PCC was markedly different. The PCC was the DMN node with maximal incoming causal influence during the first 8-10 seconds of the ABM trials, receiving input from the left AI of the SN and the left PPC of the CEN. This is also reflected in the low hubness score of the PCC (solid line in Figure 21b) and follows the predominantly anterior to posterior pattern of causal influence during this time period.

The SN nodes showed mostly incoming connections during the first 8 seconds post-task onset, compared to primarily outgoing connections during the rest of the trial. The early incoming connections were predominantly from the vmPFC, adding further evidence for vmPFC-driven memory processes during the early period of ABM trials. The CEN nodes were sparsely connected to other nodes during the first half of the ABM trials, however, showed increased PPC-driven causal influence during the latter half of the ABM trials, potentially indicative of the involvement of these nodes in maintenance of ABM.

These causal patterns were further investigated using the extended set of tri-network nodes, revealing that vmPFC remained the dominant DMN hub with maximal causal outflow during the first half of the ABM trials, and was followed by extensive causal outflow from the amPFC towards the end of the ABM trials (solid lines in Figure 20e). The causal influence from these mPFC nodes was also accompanied by high causal outflow from the right HC node throughout the ABM trials. Additionally, the newly added MTL nodes also received strong incoming connections from other DMN and SN nodes during the first 10 seconds of ABM trials (Figure 18). This is seen in Figure 20e, in the form of negative net causal flow into these nodes, and in Figure 21e, as high hubness scores for bilateral HC, pHC and RSC nodes during this time period. Furthermore, the PCC is found to be heavily connected to other tri-network nodes during the same time period, resulting in hub-like behaviour shortly after ABM task onset (Figure 21e). These results indicate highly connected hippocampal and parahippocampal hubs, that work together with a PCC hub and a mPFC hub during the first half of the ABM task, switching to a predominantly mPFC hub during the second half of the ABM trials.

The SN nodes showed numerous connections to other DMN and CEN nodes during the first half of the ABM trials, followed by a widespread increase in the hub-like properties of all SN nodes during the latter half of the ABM trials (solid lines in Figure 21d). This included the newly added PI nodes and coincided with the increase in PI activity and the second peak in MTL activity observed in Figure 11c and b, respectively. These findings might represent extensive PI-mediated processing throughout the ABM trials, and show its collaborative integration with other SN nodes.

Lastly, the addition of the MTL and PI nodes to the tri-network model led to an increase in hub-like behaviour within CEN nodes, with right PPC and dlPFC showing increased hubness score during the first half of the ABM trials, followed by a second peak in the hub-like properties of the right PPC during the latter half of the ABM trials (solid lines in Figure 21f). The continued involvement of these posterior parietal CEN nodes throughout the ABM trials might indicate a multi-functional role of these nodes in working memory and autobiographical memory recall.

Collectively, the MVGC results support an mPFC-based hub involved in autobiographical memory retrieval, that dynamically interacts with MTL nodes and other hubs anchored within SN nodes, and the regions within the parietal cortex, such as the PPC, RSC and PCC.

##### WM Trials

Similar to the ABM trials, PCC of the DMN received significant causal input from the left AI of the SN during the first 8-10 seconds of the task trials, corresponding with an increase in PCC activity during this time (Figure 9e), potentially representative of common memory-linked processes required during both ABM and WM trials.

However, contrary to the ABM trials, all SN nodes of bilateral AI and dACC exerted extensive causal influence on left CEN structures, such as PPC and dlPFC, and frontal DMN nodes such as amPFC and vmPFC during the first half of the WM task. This is reflected in the high net causal outflow from SN nodes, seen in Figure 20a (dotted lines), and high hubness score of SN nodes during this time period (dotted lines in Figure 21a). Combined with the subsequent decrease in amPFC activity after the first 8 seconds of the WM trials (Figure 9e), such causal patterns might represent the inhibition of DMN nodes, while the CEN nodes were recruited for executive processing.

Furthermore, in the first half of the WM trials, the CEN nodes of left dlPFC and left PPC received causal input from bilateral AI and dACC of the SN respectively. This was followed by extensive outgoing causal influence from bilateral PPC and right dlPFC during the latter half of the WM trials, leading to an increase in its net causal outflow and hub-like behaviour shown using dotted lines in Figures 20c and 21c, respectively. This might be representative of working memory processes mediated by the PPC and dlPFC nodes of the CEN, reaching its peak during the latter half of the WM trials.

Reanalyzing these causal patterns with the extended set of tri-network nodes revealed some finer details in the dynamic causality of the tri-network nodes. Similar to the ABM trials, the RSC was found to be a major hub in the first half of the WM trials (dotted line in Figure 21e), however, unlike the ABM trials, this was the result of primarily outgoing causality from the RSC to other tri-network nodes, as seen in Figures 19 and 20e. This was also accompanied by some hub-like behaviour from right HC and the mPFC nodes during this time period. This was followed by extensive incoming causal connections to the PreC, which in turn connected with other MTL nodes and posterior parietal CEN nodes during the second half of the WM trials, seen in Figures 19 and 20e. This led to an increase in the hub-like behaviour of the PreC, left HC and left pHC, potentially indicating a shift from RSC-centric processing early in WM trials, to more PreC-centric processing within the DMN in the latter half of the WM trials.

The newly added PI nodes also showed extensive hub-like behaviour shortly after WM task onset, alongside the bilateral AI and dACC nodes, that continued throughout the WM trials (dotted line in Figure 21d). High causal outflow was observed from these PI nodes to CEN nodes such as the dlPFC, and DMN nodes including the MTL and mPFC nodes, seen in Figure 19. These causal properties of the PI nodes added to the previously observed patterns of task-linked SN influence on DMN and CEN nodes and might warrant its inclusion in the standard set of nodes used to define the SN within the tri-network model.

Lastly, the causal patterns of the CEN nodes using the extended set of tri-network nodes were similar to that using the abridged set of tri-network nodes, with the right PPC and dlPFC showing extensive causal outflow during the first half of the WM trials, switching to mostly incoming causal links in the second half. In contrast, the left PPC and dlPFC displayed mostly incoming causal connections in the first half of the WM task, followed by some outgoing connections in the latter half of the WM trials (dotted lines in Figure 20f). This resulted in high left dlPFC hubness score shortly after WM task-onset, followed by an increase in the hub-like properties of the remaining CEN nodes during the last 15 seconds of the WM trials, coinciding with a similar increase in hub-like properties of the PreC and left HC and pHC nodes, as seen in Figures 21f and e. These results show significant functional lateralization of working memory processing within CEN nodes, and the dynamically shifting hub-like properties within the CEN nodes, which coincides with that of other DMN and MTL nodes during the second half of the WM trials.

In conclusion, unlike the ABM trials, bilateral AI and PI showed hub-like causal properties indicative of some DMN node suppression while recruiting CEN nodes for working memory processes shortly after WM task-onset, accompanied by a left dlPFC-centric hub and RSC-centric hub. This transitioned to a highly connected network structure during the latter half of the WM trials, with some HC, pHC, mPFC, PreC and all CEN nodes displaying hub-like properties, potentially indicative of working memory maintenance processes. The results also support lateralization of WM processes between left and right CEN nodes.

## 4 Discussion

The triple network model of Menon (2011) postulates that network switching between the default mode network (DMN) and central executive network (CEN), gated by the salience network (SN), is essential for everyday tasks requiring switching between internally directed and externally directed thought processes. This study adds to the growing body of evidence showing the gating function of SN while dynamically switching between two tasks specifically designed to engage these three networks. It additionally investigates the differential role of the various SN sub-networks and sub-nodes in the context of a complex switching task that utilized an autobiographical memory recall task and a 2-back working memory task to activate the DMN and CEN respectively.

The data-driven analyses performed to characterize spatial patterns of tri-network activity during the ABM and WM trials (study aim 1) identified a set of brain regions and sub-networks that are known to be associated with a wide range of functions necessary for adequate task performance, as discussed in the following paragraphs.

Autobiographical memory (ABM) retrieval is a complex multiphasic process with an initial memory instantiation/retrieval phase that is subserved by the right hippocampus, right/medial prefrontal cortices and retrosplenial cortex, followed by an elaboration phase that is found to be associated with increased activity in the left prefrontal cortex, precuneus and visual cortex (Daselaar *et al*., 2008). Each of these phases can take upto 12 seconds (Daselaar *et al*., 2008), with memory retrieval being faster for general autobiographical memories compared to specific ones (Addis *et al*., 2004). Furthermore, according to Kim (2012)‘s dual-subsystem model of the DMN, cortical midline nodes such as amPFC and PCC are involved in the self-related processing aspects of autobiographical memory (ABM) recall, while the parieto-temporal nodes, including inferior parietal lobule and medial/lateral temporal cortices, are associated with the memory retrieval aspects of the ABM task. The precuneus is also implicated in ABM retrieval (Svoboda *et al*., 2006; Addis *et al*., 2004). More specifically, the BBB model postulates a central role for the precuneus in representing the products of memory retrieval and mental imagery (Byrne *et al*., 2007). Our MVPA results (Figure 4 and Table 3) are in agreement with these findings showing activation of the medio-temporal subnetwork of the DMN, including the inferior parietal lobule of the supramarginal gyrus (SMG) and the PCC, during autobiographical memory recall (Andrews-Hanna *et al*., 2014) trials. The involvement of the parietal cortical structures such as bilateral anterior supramarginal gyri (aSMG) in autobiographical memory recall align with its involvement in self-referential processing tasks vs non self-referential processing tasks (Axelrod *et al*., 2017), and redirecting attention on internal representations (Sestieri *et al*., 2011). Furthermore, the observed ABM-linked activation in parietal opercular structures show involvement of secondary somatosensory areas during autobiographical memory recall, providing evidence for distributed constructive processes during autobiographical memory recall.

On the other hand, the N-back task is a popular working memory task that relies on the multi-component executive process of temporal coding (Collette and Van der Linden, 2002; Smith and Jonides, 1997), involving frontal brain regions such as the frontal poles, dlPFC, vlPFC, and latero-medial premotor cortices, in addition to posterior parietal regions such as medio-lateral posterior parietal cortices, dorsal cingulate (Owen *et al*., 2005; Yaple *et al*., 2019), and the precuneus (Yaple *et al*., 2019). The observed activation of the dlPFC in the MidFG (Figure 4 and Table 3) during the working memory trials indicates activity of this core fronto-parietal CEN subsystem (Yaple *et al*., 2019). Furthermore, the engagement of supplementary motor areas might indicate the involvement of sub-vocal rehearsal structures (Smith and Jonides, 1997; Fegen *et al*., 2015) for maintaining the presented words in working memory through articulatory rehearsal processes (Vallar and Pagano, 2002).

The ICA analyses identified data-driven differences in activity, finding two functionally distinct sub-networks within each of the three networks (Figure 5). These sub-networks significantly overlapped with the networks presented in Shirer *et al*. (2012) and showed distinct activation patterns between the ABM and WM trials, indicating functionally distinct activation of various nodes within each of the networks to accomplish the complex ABM and WM tasks.

Characterization of the temporal and causal patterns of tri-network (study aims 1 and 2 respectively) yielded five key findings, which are discussed below.

### 4.1 Different DMN and CEN nodes and sub-networks activated during both task conditions

Both DMN and CEN were observed to be active during the ABM and WM trials, although different nodes and sub-networks of the networks were predominantly active during each task (Figures 6 and 8).

DMN activity during ABM trials was expected due to its key involvement in self-related processing and memory retrieval (Spreng *et al*., 2010), however, contrary to expectations, the CEN was also observed to be active during the ABM trials, as seen in Figures 6a and 7a, where the activity of the CEN network and its sub-networks increased during the latter half of the ABM trials. Figure 9a shows that the observed increase in CEN activity was due to an underlying increase in dlPFC activity which might have been due to the working memory needed to maintain the retrieved memory throughout the trial duration, once it had been recalled. This increase in dlPFC activity began around 8-12 seconds post-task onset, aligning with the time required to fully retrieve an ABM (Addis *et al*., 2004; Daselaar *et al*., 2008). This could also have been due to the involvement of executive processing during autobiographical memory recall (Unsworth *et al*., 2012) used for searching autobiographical information. Such a pattern of DMN recruitment at the onset of a DMN-linked task, followed by CEN recruitment is also seen in creative idea production (Beaty *et al*., 2015).

Furthermore, the hippocampus and associated medial temporal lobe structures (MTL) co-activated with the aforementioned peak in dlPFC and PPC activity (Figure 11a and b). This was accompanied by an increase in hub-like behaviour within the HC, pHC and RSC nodes shortly after ABM task-onset, followed by a second increase in hub-like activity of the right PPC and some MTL nodes during the latter half of the ABM trials (Figure 21e and f). These results are consistent with the BBB model that implicates the hippocampus and associated MTL structures in the retrieval and reconstruction of details associated with a particular autobiographical memory, followed by its representation within parietal lobe structures, forming mental imagery representations within the “parietal window” that enables conscious access to these products of memory retrieval (Byrne *et al*., 2007).

During WM trials, the DMN was also found to be active in addition to the expected CEN activity. More specifically, the dorsal DMN sub-network was found to be active during the first half of the WM trials, compared to the ventral DMN being predominantly active during the ABM trials (Figure 7e). This difference can be attributed to the higher activity of the amPFC (a node of the dorsal DMN) during the WM trials, in contrast to the ABM trials (Figure 9e). Another contributing factor could have been the participation of posterior parietal nodes of the dorsal DMN sub-network (Figure 5), which are also associated with n-back working memory tasks (Owen *et al*., 2005). These posterior parietal nodes also showed increased hub-like behaviour towards the latter half of the WM trials (Figure 21f), a causal pattern that was also observed within the precuneus and some other MTL nodes (Figure 21e), consistent with the role of these parietal structures in maintaining mental images of items in working memory (Byrne *et al*., 2007).

Interestingly, the PCC node of the DMN also showed increased activity during the initial phase of the WM task, followed by a large decrease in activity when DMN nodes increased their activity later in the WM trials (Figure 9e). PCC activity was also observed to increase during the initial 8-10 seconds of the ABM trials, followed by a subsequent decrease in activity, however, not decreasing to the extent seen during the WM trials. Although seemingly contrary to the expected inactivation of PCC during WM, the observed pattern is consistent with the role of PCC in self-related processing, while also participating in cognitive control tasks under high task load (Leech *et al*., 2011).

### 4.2 The SN recruited the task-appropriate network by synchronizing its activity with the desired network

Despite both DMN and CEN being active during both conditions, global SN activity was found to correlate with the task-appropriate network, and anti-correlate with the “task-opposite” network. Figure 8 shows that SN activity was correlated with the early peak in DMN activity during ABM trials and the late peak in CEN activity during WM trials and that it was anti-correlated with the later peak in CEN activity during ABM trials and the early peak in DMN activity during WM trials. The peaks in global SN activity were also observed to lead the corresponding peaks in DMN or CEN activity by 2-4 seconds (Figure 8). These findings, combined with the observed causal patterns of SN nodes driving the DMN and CEN nodes (Figure 15), indicate that the SN might have been gating the activity of the corresponding network by synchronizing its activity with that of the task-appropriate network. Interestingly, this pattern of network co-activation is similar to that observed within the dual-network model of cognitive control, wherein the salience network (cingulo-opercular network) is observed to co-activate with the central executive network (fronto-parietal network) during executive task performance (Dosenbach *et al*., 2008). The dual-network model ascribes this pattern of co-activation to the set-maintenance role of the SN, which when combined with its role in salience assignment (Menon, 2011) could explain its gating function within the tri-network model. In this paradigm, the SN would use key salience signals to identify the task-appropriate network based on the interoceptive or exteroceptive nature of the task demands, and then co-activate with the identified network to stably maintain task sets during task performance. This effectively selects the task-appropriate network by providing stable set-maintenance to the network deemed appropriate for the task at hand, following the general principle of ‘connection through coherence’ (Fries, 2015), albeit in the context of network connectivity.

### 4.3 Different sub-networks of the SN showed different causal properties

Data driven ICA analysis revealed two distinct sub-networks that have significant overlap with SN nodes (Figure 5), forming anterior and posterior salience sub-networks, similar to those described by Shirer *et al*. (2012). The anterior SN seemed to co-activate with the task-appropriate network in both ABM and WM trials, however, posterior SN seemed to correlate with bilateral CEN and dorsal DMN activity (Figure 7). While the anterior SN causally influenced most of the other sub-networks, owing to its hub-like properties, the posterior SN network causally inhibited ventral DMN activity during the latter half of the ABM trials, while causally stimulating dorsal DMN activity during the first half of the WM trials (Figure 13).

Posterior SN was also closely linked to the LCEN activity, providing causal input during both WM and ABM trials. This could be owing to its role in maintaining a representation of the passage of time (Wittmann *et al*., 2010), which could be important for successfully performing an N-back task by keeping track of the word seen 2 words ago. Posterior SN deactivation has also been observed in the case of 2-back verbal working memory (Sweet *et al*., 2008). This combination of posterior SN activation, owing to its time-keeping role, and its deactivation observed in n-back working memory tasks might explain the activation-inactivation pattern of posterior SN activity observed in Figure 7c and f.

The posterior insula (PI) is one of the major constituent nodes of the posterior SN subnetwork, as previously described (Shirer *et al*., 2012) and seen in Figure 5. However, its role in network switching has been relatively understudied compared to that of the AI (Menon and Uddin, 2010) and dACC (Crottaz-Herbette and Menon, 2006). The high hubness score of the PI and extensive integration with other SN, DMN and CEN nodes observed in this study (Figures 18, 19, and 21d) indicate the dynamic role played by the PI in autobiographical memory retrieval and working memory processes. The PI is well positioned to integrate various aspects of salient information and guide network switching, given its multi-sensory inputs (Björnsdotter *et al*., 2009), and its connectivity with the emotionally salient ventral AI, and the cognitively salient dorsal AI (Davidovic *et al*., 2019). Dysfunctional PI activity has also been observed with abnormal cognitive behaviour, as seen in autism spectrum disorders (Ebisch *et al*., 2011) and PTSD (Nicholson *et al*., 2020).

### 4.4 Distinct causal roles of the right AI and the left AI

As has been extensively discussed in the literature, the right AI is a key hub region that modulates network switching by recruiting the CEN and inhibiting the DMN during externally directed tasks (Sridharan *et al*., 2008), as also seen in this study by the patterns of rAI to amPFC/vmPFC causality (Figure 20, panel a and d, and Figure 17). However, this study also found that the right AI was not the primary causal outflow hub during self-directed processing, and that the left AI alongside the dACC and vmPFC played a much larger role in recruiting other DMN nodes when required in the ABM trials (Figure 20, panel a and d, and Figure 16).

In light of the results from sections 4.3 and 4.4, the bilateral insula seems to form a set of key nodes that modulate inter-network connectivity in a task-linked manner, and has been also observed to couple with the DMN at the onset of creative idea production (Beaty *et al*., 2015).

### 4.5 Left CEN nodes formed a separate cluster (from Bilateral CEN) that was uniquely activated for each of the two tasks

The data-driven ICA analyses revealed that left CEN nodes engaged in distinct connectivity patterns, compared to the rest of the CEN (Figure 14), and was hence identified as an independent spatiotemporal component (Figure 5) by the data-driven group-ICA algorithm. This was further supported by the ROI-to-ROI analyses revealing primarily left CEN nodes receiving incoming connections during the onset of WM trials (Figures 15 and 20c/f). Left lateralization has been extensively documented in some key working memory sub-systems, most notably the subvocal rehearsal sub-systems of the phonological loop (Smith and Jonides, 1997; Fegen *et al*., 2015). This might also be the underlying reason for the observed increase in hubness score of the left dlPFC during the first half of the WM trials (Figure 21f), and the subsequent increase in activity of the left CEN sub-network during the latter half of the WM trials (Figure 7).

These findings consolidate a normative view of network dynamics within Menon‘s tri-network model, that had originally emerged from a large body of clinical research, suggesting that many psychopathologies can be viewed as a dysregulation of these three key networks. For example, an overly dominant DMN is associated with maladaptive rumination in depression (Hamilton *et al*., 2011), while dysregulated SN nodes lead to aberrant network switching in schizophrenia (Moran *et al*., 2013), and Post-traumatic stress disorder (PTSD) (Lanius *et al*., 2015). PTSD is an especially relevant example with patients suffering from improper recruitment of the DMN during a working memory task (Daniels, 2010), while their DMN was under-recruited during an autobiographical memory task (St. Jacques and De Brigard, 2015). Given the critical role of the SN and its sub-networks in healthy network-switching, such PTSD-linked network dysfunctions might be due to an aberrant SN causing disruptions in normal salience detection (Rabellino *et al*., 2015). This might lead to a breakdown of the SN-linked co-activation dynamics described in section 4.2, which can provide a mechanistic explanation of how abnormal recruitment of task-opposite networks can lead to poor task performance in patients with PTSD. For example, the abnormal recruitment of DMN during a working memory task observed in PTSD patients (Daniels, 2010), is also associated with an absence of correlated activity between SN and CEN. This absence of SN-linked stable set-maintenance might result in the rapid decay of the working memory items maintained by the CEN, ultimately resulting in poor working memory performance in patients with PTSD. Such improved understanding of the mechanisms underlying healthy and dysfunctional SN-gated switching between DMN and CEN could inform the development of specialized treatment plans that directly target the dysfunctional tri-network dynamics (Lanius *et al*., 2015) in patients with various psychopathologies.

This study used MVGC analysis to probe the patterns of causal influence between the SN, CEN and DMN. However, some considerations should be kept in mind when using MVGC analysis with BOLD fMRI signals due to the poor time resolution of the BOLD signal, reliance of the BOLD signal on the haemodynamic response, and the susceptibility of GC estimates to poor signal-to-noise ratio (SNR) (Ramsey *et al*., 2010). Despite these concerns, GC estimates have been found to be robust in the context of haemodynamic response function confounds (Schippers and Keysers, 2011), and its multivariate version is found to appropriately represent the connectivity patterns in low density networks (Duggento *et al*., 2018). The impact of the other factors can be minimized by using sliding windows to maximize signal stationarity, performing “vertical regression” across trials that are expected to have similar stochastic generative processes, and comparing the MVGC estimates of the signal with those of the time-permuted surrogate signals to minimize spurious connectivity estimates, as performed here.

One caveat to the ROI analyses performed in this study is that the network membership assigned to inherently multi-functional nodes such as vmPFC and PPC is based on the definitions used by the current work surrounding this tri-network model (Menon, 2011). However, as evident from the BBB model, there is a complex interplay between fronto-parietal areas in encoding, maintaining and manipulating the products of memory retrieval (Byrne *et al*., 2007). For example, in addition to its role within the DMN, vmPFC is also an integral component of reward processing circuits and can be heavily involved in some executive tasks (Gerlach *et al*., 2013). The PPC, similarly, has a broader role in representing mental imagery, which is important for both working memory and maintaining the retrieved components of autobiographical memory (Byrne *et al*., 2007). This is supported by our results showing activation of these structures during the ABM trials (Figure 4) and WM trials (Figure 9). Thus nodes such as vmPFC and PPC can be expected to dynamically shift network membership, as a function of dynamic cognitive processing, and should be considered as cross-network nodes (Pedersen *et al*., 2018). As such, these multifunctional nodes are considered within the context of the tri-network model in this study, however, future studies could look into a dynamic version of the tri-network model with flexible node-membership.

In conclusion, results from the current study add to the growing body of evidence showing the complex interplay of CEN, DMN and SN nodes and sub-networks required for adequate task-switching. Furthermore, the discovery that the SN co-activates with the task-relevant network provides mechanistic insight into SN-mediated network selection in the context of explicit tasks. Finally, the results from this study indicate active involvement of the posterior insula and some MTL nodes in task-linked functions of the SN and DMN, warranting their inclusion as network nodes in future studies of the tri-network model.

## 5 Acknowledgements

This research was supported by a Discovery Grant from the Natural Sciences and Engineering Research Council (NSERC) of Canada to SB, an Alexander Graham Bell Canada Graduate Scholarship (CGS-D) from NSERC Canada to SBS, a Canadian Institutes of Health Research (CIHR) grant to MCM and a NSERC Canada grant to JH. MCM is also supported by the Homewood Chair in Mental Health and Trauma at McMaster University.

## Declarations of Interest

None

## Notes

### Competing Interest Statement

The authors have declared no competing interest.

